# The RNA-binding protein RSRC2 promotes mitotic fidelity by interacting with the lncRNA *C1QTNF1-AS1*

**DOI:** 10.1101/2025.01.21.634161

**Authors:** Kaliya Georgieva, Alice O Coomer, Parnia Babaei, Giulia Guiducci, Martin Dodel, Eleni Maniati, Anisha Thind, Sneha Krishnamurthy, Anna Nawrocka, Sam Wallis, Alena Shkumatava, Jun Wang, Faraz K Mardakheh, Lovorka Stojic

## Abstract

Mitotic fidelity depends on proper chromosome alignment at the spindle equator, a process known as chromosome congression, driven by well-established protein networks. Whereas RNA-binding proteins and noncoding RNAs have been implicated in cell division, their interplay during this process remains unknown. Here, we discover that RSRC2, an arginine/serine-rich RNA-binding protein, plays an essential role in cell division by interacting with the long non-coding RNA *C1QTNF1-AS1*. The loss of either RSRC2 or *C1QTNF1-AS1* results in defects in chromosome congression and mitotic progression. We show that RSRC2 interacts with distinct sets of proteins involved in splicing and centrosome biogenesis, contributing to the fidelity of cell division through two different mechanisms: one linked to the splicing of mitotic regulators and the other by localising to mitotic centrosomes for which the interaction with the *C1QTNF1-AS1* RNA is required. Our study uncovers RSRC2 as a new regulator of cell division and illustrates how RNA-protein complexes promote error-free mitosis.

## Introduction

Chromosome congression, the process of aligning chromosomes at the spindle equator, is a prerequisite for faithful chromosome segregation during cell division^1^. In human cells, this process lasts 15-20 minutes and facilitates the formation of the metaphase plate, which is spatially and temporally coordinated with the assembly of the mitotic spindle. The mitotic spindle apparatus is a microtubule-based structure that mediates the kinetochore-microtubule interactions required for chromosome movement during cell division. Multiple mechanisms involving more than 100 characterised proteins, from kinases and phosphatases to kinetochores, centrosomes and motor proteins, have been implicated in chromosome congression^2^. Any perturbations in these processes leading to alterations in microtubule dynamics or kinetochore function will affect chromosome congression and lead to chromosomal instability, a hallmark of cancer^3^.

In addition to well-established protein networks, emerging evidence suggests the role of various types of RNAs, from protein-coding mRNAs^4, 5^ to non-coding RNAs (ncRNAs) ^6–11^ and RNA-binding proteins (RBPs)^12, 13^ in the structural and functional integrity of the mitotic spindle apparatus^12^. Multiple RBPs contribute to the control of cell division through post-transcriptional gene regulation^14^ either directly by localising to the mitotic spindle^15, 16^ or indirectly by regulating the splicing of pre-mRNAs required for cell division^17–19^. For example, an RNA splicing factor SON^17, 18^ is not present on the mitotic spindle; however, its depletion leads to mitotic defects caused by the mis-splicing of many mitotic pre-mRNAs. Other splicing regulators have a direct role in mitosis by interacting with centrosomes^15^, spindle microtubules^20^ and/or the Ndc80 complex, which is essential for kinetochore-microtubule attachment ^21^. In another study, the PRP19 splicing complex was shown to have both direct and indirect roles in cell division^22, 23^. Furthermore, several RNAi genome-wide screens conducted in human cells demonstrate that the depletion of splicing factors leads to a variety of mitotic defects^24–26^, highlighting their importance in regulating cell division. Indeed, the recent mapping of proteogenomic interactions shows that spliceosome components play a significant role in the centrosome-cilia network^27^. Consequently, many splicing factors exhibit moonlighting roles during mitosis, independent of RNA splicing^28^. However, how the regulation of the dual roles of these moonlighting proteins is controlled, both temporally and spatially, remains unclear.

Long noncoding RNAs (lncRNAs) have also been implicated in the control of cell division^29^. LncRNAs are a heterogeneous class of RNA molecules localised in the cytoplasm and the nucleus^30^. Regardless of their sub-cellular localisation, lncRNAs interact with RBPs. RBPs have a key role in driving the functions of lncRNAs with binding specificity typically determined by RNA motifs or structural elements present within those lncRNAs^31^. Despite some mechanistic studies where lncRNAs were implicated in mitotic progression^7, 8^, very little is known about whether (lnc)RNA-mediated regulation of RBPs is involved in the fidelity of cell division.

Herein, we report that a lncRNA *C1QTNF1-AS1* regulates cell division by interacting with RSRC2, an RBP whose function is largely unknown. We show that depletion of either *C1QTNF1-AS1* or RSRC2 results in similar phenotypic outcomes, causing mitotic delay and chromosome congression defects. Loss of RSRC2, but not *C1QTNF1-AS1*, results in the mis-splicing of transcripts enriched for cell cycle functions, and RSRC2 protein-protein interactions include centrosome and splicing proteins as its top interactors. Furthermore, co-depletion of RSRC2 and *C1QTNF1-AS1* did not have additive effects, suggesting that they both act in the same pathway during cell division. We also demonstrate that RSRC2 recruitment to mitotic centrosomes is regulated by *C1QTNF1-AS1*. Thus, our work uncovered that RSRC2 plays dual roles in cell division. One that is independent of *C1QTNF1-AS1* and regulates the splicing of several mitotic proteins, and another that has mitotic function and is dependent on *C1QTNF1-AS1*. Based on these results, we propose that *C1QTNF1-AS1* acts as a regulator of RSRC2’s mitotic but not splicing activity to safeguard mitotic fidelity. Our findings support the rapidly emerging evidence that ncRNAs serve as rheostats of protein functions to promote error-free mitosis.

## Results

### *C1QTNF1-AS1* lncRNA interacts with the uncharacterised RNA-binding protein RSRC2

We previously identified *C1QTNF1-AS1* as a lncRNA crucial for mitotic progression in HeLa cells^7^. *C1QTNF1-AS1*, as annotated in GENCODE, is antisense to a protein-coding gene *C1QTNF1/CTRP1* and conserved between mouse and human (Fig. 1A). To assess the role of *C1QTNF1-AS1* in cell division, we used chromosomally stable and near diploid human cells hTERT-RPE1 (hTERT-immortalised retinal pigment epithelium, hereafter referred to as RPE1) and HCT116 cells (colon carcinoma), both widely used to study cell division. Consistent with the *C1QTNF1-AS1* localisation in HeLa cells^7^, single-molecule RNA fluorescence in situ hybridisation (smRNA FISH) showed that *C1QTNF1-AS1* is predominantly localised in the nucleus of HCT116 and RPE1 cells (Fig. 1B; Supp. Fig. 1A). Three LNA gapmers^32^ targeting different regions of *C1QTNF1-AS1* led to the downregulation of the *C1QTNF1-AS1* transcript levels and its RNA FISH signal (Fig. 1C, Supp. Fig. 1B). To test whether, in addition to interphase DNA, *C1QTNF1-AS1* localises to the mitotic spindle, we combined smRNA FISH with immunofluorescence (IF), staining the cells for α- and y-tubulin to visualise the microtubules and centrosomes of the mitotic spindle, respectively. We observed that *C1QTNF1-AS1* is not present on the mitotic spindle (Supp. Fig. 1C), consistent with previous findings indicating that lncRNAs lack mitotic bookmarking functions^33^.

**Figure 1.**
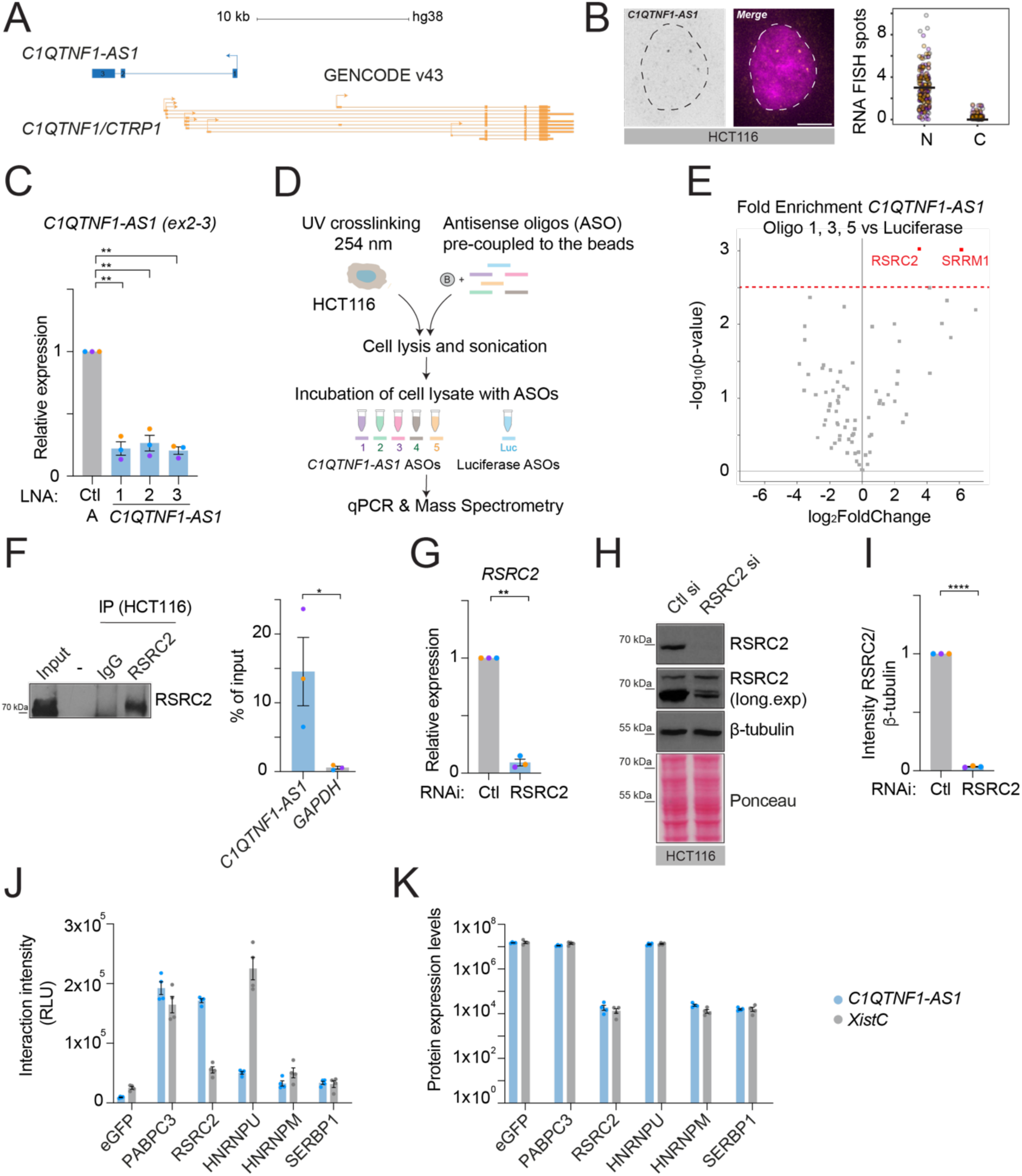
*C1QNTF1-AS1* interacts with RSRC2, an RNA-binding protein of unknown function. (A) Schematic representation of the *C1QTNF1-AS1* and *C1QTNF1/CTRP1* genomic landscape (*C1QTNF1-AS1* annotated as NR_040018/NR_040019 in RefSeq; Gencode gene *ENSG00000265096*; chr17:79019209-79027601, hg38). (B) Maximum intensity projections of representative images of *C1QTNF1-AS1* exon smRNA FISH in HCT116 showing its nuclear localisation. Nuclei were stained with 4,6-diamidino-2-phenylindole (DAPI, magenta) and outlined with a dashed circle. Right panel: quantification of total transcript in the nucleus (N) and cytoplasm (C), solid line represents the mean. N = 3 (n_(HCT116) =_398). (C) Expression levels of *C1QTNF1-AS1* in HCT116 cells following depletion with three LNA gapmers targeting either exon 3 (LNA 1) or first intron of *C1QTNF1-AS1* (LNA 2, 3), as measured by qPCR. Primers spanning mature (ex2-3) *C1QTNF1-AS1* were used. Results are presented relative to negative control (Ctl A) LNA. N = 3. (D) Schematic representation of workflow for the antisense oligonucleotide (ASO) pulldown of *C1QTNF1-AS1* in HCT116 cells. Five different ASOs targeting different regions of the *C1QTNF1-AS1* locus were used, with luciferase (Luc) ASOs as a negative control. Pulldown efficacy was assessed by qPCR (Supp. Fig. 1D) and proteins were identified using LC-MS analysis (see Methods). (E) The volcano plot highlighting proteins enriched in *C1QTNF1-AS1* pulldown using ASO 1, 3 and 5 versus Luc. Significant *C1QTNF1-AS1* protein interactors are highlighted in red (FDR 5%). (F) RIP-qPCR from HCT116 extracts. Left panel: Western blot of RSRC2 in the input and IP samples to show RSRC2 IP efficiency compared to IgG. Right panel: RIP-qPCR showing association of RSRC2 with *C1QTNF1-AS1* transcript. *GAPDH* was used as negative control RNA for RSRC2 RIP. RIP enrichments are presented as % of input RNA (normalized to IgG). N = 3. (G) Relative expression of *RSRC2* in HCT116 cells following siRNA-mediated depletion of RSRC2, as measured by qPCR. Results are presented relative to control siRNA (Ctl). N = 3. (H) Representative western blot showing RSRC2 protein expression in HCT116 cells following siRNA-mediated depletion of RSRC2. β-tubulin and Ponceau staining were used as loading controls. (I) Densitometric analysis of RSRC2 levels from panel (H) relative to control siRNA (Ctl). N = 3. (J) Interaction intensities between *C1QTNF1-AS1* and the indicated proteins show that *C1QTNF1-AS1* interacts with RSRC2. In-cell interactions were measured in an incPRINT experiment where MS2-tagged *C1QTNF1-AS1* RNA was co-expressed with a set of FLAG-tagged proteins in HEK293T cells harbouring a luciferase detector fused to the MS2 coat protein (MS2CP). Upon the formation of FLAG-protein-RNA-MS2-MS2CP ternary complexes, RNA-protein interactions were measured by luciferase activity^35^. eGFP (enhanced green fluorescent protein) was used as a negative control and PABPC3 (a polyadenylated RNA binding protein) was used to control for RNA expression. *Xist(C)-* MS2 vector was used alongside *C1QTNF1AS1-*MS2 as a positive control for RNA-protein interactions. RLU are relative light unit. N = 4. (K) Protein expression levels were estimated from horseradish peroxidase ELISA of FLAG-tagged proteins in the same experiment as in J. Error bars in all panels are shown as mean ± S.E.M. Scale bar, 5 μm. N = number of cells analysed. An unpaired t-test with Welch’s correction was applied in panels (C), (G) and (I). Unpaired t-test was used in panel (F).

As the first step toward defining *C1QTNF1-AS1’s* cellular function and molecular mechanism of action, we set out to identify proteins *C1QTNF1-AS1* interacts with. We conducted antisense oligonucleotide RNA pulldown coupled with mass spectrometry (MS) (Fig. 1D) using three *C1QTNF1-AS1* oligonucleotides (1, 3, 5) that specifically purified *C1QTNF1-AS1* from UV-crosslinked HCT116 cells (Supp. Fig. 1D). Two proteins, RSRC2 and SRRM1, previously identified in a database of RBPs^34^ were significantly enriched relative to the Luciferase (Luc) RNA sequence control sample (Fig. 1E; Supp. Table 1). We focused on RSRC2 since its depletion has been reported to cause mitotic delay in the Mitotic Cell Atlas database^25^. To validate the interaction between *C1QTNF1-AS1* and RSRC2, we employed RNA immunoprecipitation (RIP) and in-cell protein-RNA interaction (incPRINT)^35^. RIP-qPCR experiments in HCT116 using the antibody against RSRC2 detected enrichment of *C1QTNF1-AS1* compared to *GAPDH* (Fig. 1F). The specificity of the RSRC2 antibody was tested by depleting RSRC2 with a pool of four siRNAs and confirming the knockdown efficiency by qPCR and Western blot (Fig. 1G-I). Furthermore, we confirmed the *C1QTNF1-AS1*-RSRC2 interaction using the incPRINT assay^35^. Analysis of incPRINT luminescence revealed specific interactions of *C1QTNF1-AS1*-MS2 and RSRC2-FLAG but not with other FLAG-tagged proteins used as controls (Fig. 1J). We included lncRNA *Xist* (C region)-MS2 RNA as a positive control and confirmed that *Xist* interacts with hnRNPU, as reported for the endogenous transcript^36^. To control for expression levels of selected proteins, their abundance was analysed using an ELISA assay, which confirmed that the levels of the selected proteins remained unchanged across all conditions (Fig. 1K). Together, several orthogonal methods demonstrate a molecular interaction between the *C1QTNF1-AS1* lncRNA and RSRC2 protein.

### Loss of either *C1QTNF1-AS1* or RSRC2 causes defects in chromosome alignment and mitotic progression

To assess the role of *C1QTNF1-AS1* and RSRC2 in cell division, IF using a kinetochore (CREST) and microtubule (α-tubulin) was performed in cells depleted of *C1QTNF1-AS1* and RSRC2. LNA-mediated depletion of *C1QTNF1-AS1* caused chromosome congression defects in 27-36% of mitotic cells as compared to 14% in control LNA A-treated cells (Fig. 2 A, B). Since the lncRNA loci are functional at the transcript (RNA) level and/or on the transcriptional (DNA) level^30^, we employed CRISPR/Cas13 as an alternative loss-of-function method used for targeted RNA knockdown^37^. Similar to LNA-mediated depletion, Cas13-mediated depletion of *C1QTNF1-AS1* also led to chromosome congression defects, further validating the *C1QTNF1-AS1* phenotype (Fig. 2 C-E). Time-lapse imaging of HCT116 cells stably expressing GFP-tagged histone H2B (H2B-GFP) confirmed that *C1QTNF1-AS1* downregulation causes mitotic delay. Indeed, cells treated with control (Ctl A) initiated anaphase onset at 37 min, while the cells depleted of *C1QTNF1-AS1* with LNA 1, 2 and 3 gapmers initiated anaphase onset at 60, 45 and 49 min, respectively (Fig. 2F). In addition, time-lapse microscopy of *C1QTNF1-AS1*-depleted RPE1 H2B-GFP cells also led to a small but a significant delay (2-4 min) in mitotic duration (Supp. Fig. 1E, F). Given that mitotic defects can induce a p53-dependent delay in cellular proliferation and cell cycle progression in the subsequent cell cycle^38^, we analysed these processes upon *C1QTNF1-AS1* loss. No major effects on cell proliferation in HCT116 p53+/+ and HCT116 p53-/-cells (Supp. Fig. 1G), and cell cycle progression were observed (Supp. Fig. 1H, I). These data indicate that the loss of *C1QTNF1-AS1* does not lead to a p53-mediated delay in proliferation and cell cycle progression. In contrast, a gain-of-function (GOF) experiment revealed that overexpression using the expression plasmid encoding *C1QTNF1-AS1* cDNA (full) did not affect the mitotic timing compared to HCT116 cells overexpressing a scrambled (scr) *C1QTNF1-AS1* DNA sequence (Supp. Fig. 1J, K). smRNA FISH showed that subcellular localisation has not changed upon overexpression of *C1QTNF1-AS1* (full) (Supp. Fig. 1L). Based on these results, we conclude that the role of *C1QTNF1-AS1* in cell division is dependent on its RNA.

**Figure 2.**
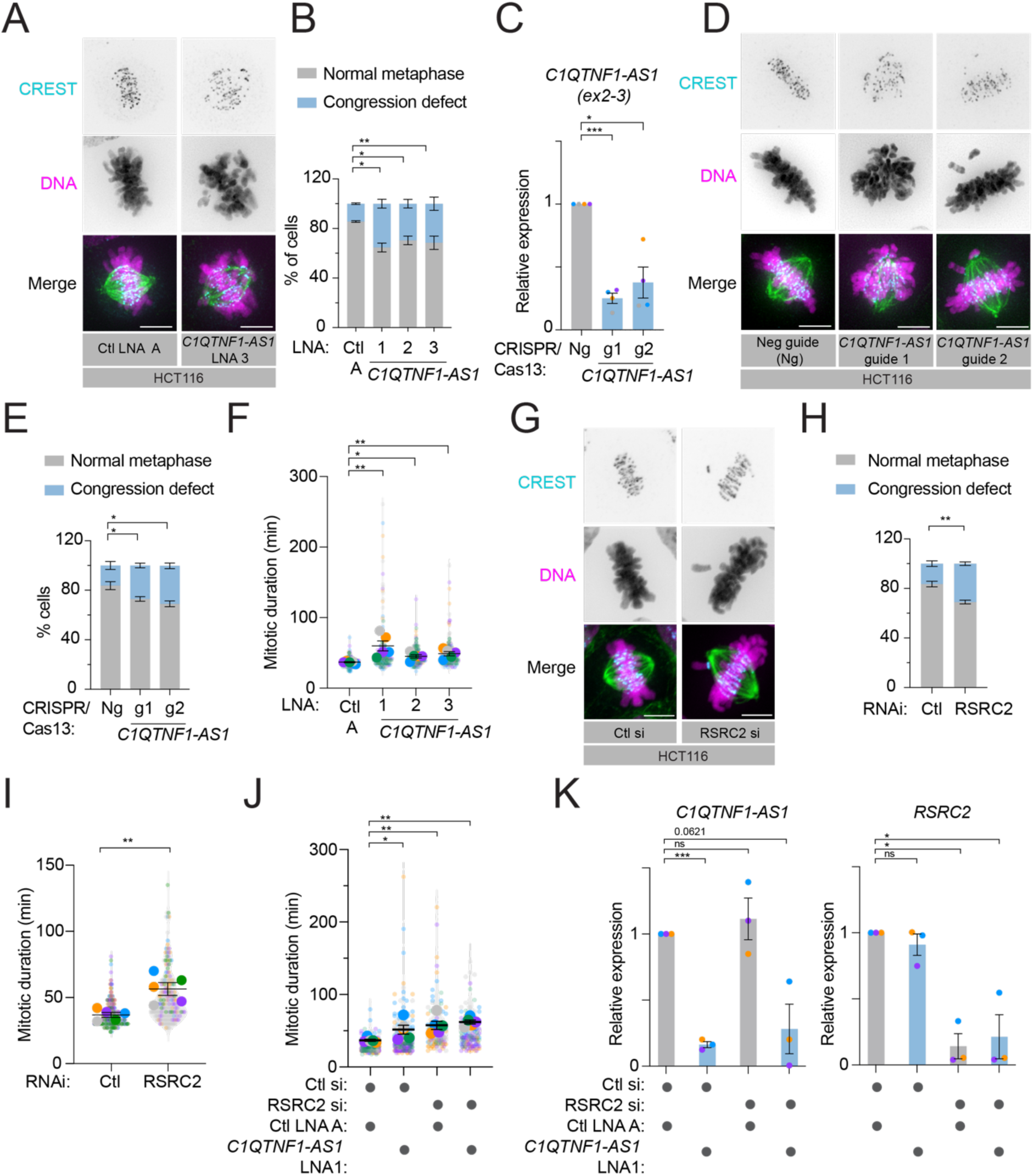
*C1QTNF1-AS1* and RSRC2 are critical for chromosome alignment and mitotic progression. (A) Representative images of metaphase cells stained for kinetochores (CREST, cyan), microtubules (α-tubulin, green) and DNA (Hoescht, magenta) after LNA-mediated depletion of *C1QTNF1-AS1* in HCT116 cells. (B) Characterisation of mitotic phenotypes in HCT116 cells after LNA-mediated depletion of *C1QTNF1-AS1*. Percentage of mitotic cells with normal metaphase plate and congression defect using the same antibodies as in panel (A). N = 3 (n_(Ctl A)=_627; n_(*C1QTNF1-AS1* LNA1)=_578; n_(*C1QTNF1-AS1* LNA2)=_560; n_(*C1QTNF1-AS1* LNA3)=_586). (C) Expression levels of *C1QTNF1-AS1* after CRISPR/Cas13-mediated depletion of *C1QTNF1-AS1* using two guide RNAs targeting *C1QTNF1-AS1* relative to the negative guide RNA (Ng), as measured by qPCR. N = 4. (D) Representative IF images of metaphase cells after CRISPR/Cas13-mediated depletion of *C1QTNF1-AS1* in HCT116 cells using the same antibodies as in panel (A). (E) Characterisation of mitotic phenotypes in HCT116 cells following CRISPR/Cas13-mediated depletion of *C1QTNF1-AS1* as in panel (D). N = 4 (n_(Ng)=_439; n_(*C1QTNF1-AS1* g2)=_471; n_(*C1QTNF1-AS1* g3)=_424). (F) Quantification of mitotic duration from time-lapse imaging microscopy following LNA-mediated depletion of *C1QTNF1-AS1* in HCT116 H2B-GFP cells. Mitotic duration was measured from nuclear envelope breakdown to anaphase onset. N = 5 (n_(Ctl A)=_180; n_(*C1QTNF1-AS1* LNA1)=_177; n_(*C1QTNF1-AS1* LNA2)=_177; n_(*C1QTNF1-AS1* LNA3)=_196). (G) Representative images of metaphase HCT116 cells stained for kinetochores (CREST, blue), microtubules (α-tubulin, green) and DNA (Hoescht, cyan) following siRNA-mediated depletion of RSRC2. (H) Characterisation of mitotic phenotypes in HCT116 cells after siRNA-mediated depletion of RSRC2. Percentage of mitotic cells with normal metaphase plate and congression defect using the same antibodies as in panel (G). N = 3 (n_(Ctl si)=_289; n_(RSRC2 si) =_271). (I) Quantification of mitotic duration from time-lapse imaging microscopy following siRNA-mediated depletion of RSRC2 in HCT116 H2B-GFP cells. N = 5 (n_(Ctl si)=_308; n_(RSRC2 si)=_192). (J) Quantification of mitotic duration from time-lapse imaging microscopy after double knockdown of *C1QTNF1-AS1* and RSRC2 in HCT116 H2B-GFP cells. N = 5 (n_(Ctl si+Ctl LNA A)=_244; n_(Ctl si+C1QTNF1-AS1 LNA1)=_232; n_(RSRC2 si+C1QTNF1-AS1 LNA1)=_115; n_(RSRC2 si+ Ctl LNA A)=_152). (K) Expression levels of *C1QTNF1-AS1* and RSRC2 in HCT116 cells after the double knockdown experiment (as in panel J) were measured by qPCR. Results are presented relative to negative controls (Ctl LNA A + Ctl si). N = 3. Error bars in all panels are shown as mean ± S.E.M. Scale bar, 5 μm. An unpaired t-test with Welch’s correction was applied in panels (B), (C), (E), (H) and (K). Mann-Whitney test was used in panels (F), (I) and (J).

Given that antisense lncRNAs such as *C1QTNF1-AS1* can regulate the expression of their overlapping sense protein-coding partner^30^, we investigated whether *C1QTNF1-AS1* affects the expression of the *C1QTNF1/CTRP1* gene. Depletion of *C1QTNF1-AS1* using LNA gapmers did not alter the levels of *C1QTNF1/CTRP1* expression in HCT116 cells, as measured by qPCR using primers detecting the total expression of three isoforms (NM_198593, NM_030968, NM_153372, also known as *C1QTNF1/CTRP1*) (Supp. Fig. 2A). This finding was further confirmed with CRISPR/Cas13 knockdown of *C1QTNF1-AS1* in HCT116 cells (Supp. Fig. 2B). Furthermore, *C1QTNF1-AS1* depletion did not affect C1QTNF1/CTRP1 protein levels (Supp. Fig. 2C, D). Since many antisense lncRNAs can regulate splicing of their sense pre-mRNA^39^, we also examined the expression of all annotated *C1QTNF1/CTRP1* splice variants and found no changes in their isoform-specific expression (Supp. Fig. 2E). Thus, we concluded that *C1QTNF1-AS1* does not affect *C1QTNF1/CTRP1* splicing. Depletion of C1QTNF1/CTRP1 using a pool of siRNA sequences (Supp. Fig. 2F-H) did not affect chromosome congression (Supp. Fig. 2I, J) and did not cause a delay in mitotic duration in RPE1 H2B-GFP cells (Supp. Fig. 2K, L). Together, these results indicate that C1QTNF1/CTRP1 is neither regulated by its neighbouring lncRNA, nor involved in the regulation of cell division.

We next sought to elucidate the function of the *C1QTNF1-AS1* protein interactor RSRC2 in cell division. Loss of RSRC2 did not show any changes in cell cycle progression (Supp. Fig. 3A, B), similar to the *C1QTNF1-AS1* depletion (Supp. Fig. 1H, I). However, RSRC2-depleted cells showed a significant increase in misaligned chromosomes, with nearly 30% of mitotic cells exhibiting chromosome congression defects compared to 16% in control siRNA-treated cells (Fig. 2 G, H). In line with these results, HCT116 H2B-GFP cells treated with control siRNA (Ctl siRNA) initiated anaphase onset at 37 min, while RSRC2-depleted cells initiated anaphase onset at 57 min (Fig. 2I). Similarly, time-lapse imaging of RPE1 H2B-GFP cells showed that RSRC2 depletion led to a mitotic delay of 4 min compared to control siRNA-treated cells (Supp. Fig 3C, D). Thus, the RSRC2 knockdown phenocopies the *C1QTNF1-AS1* LOF mitotic phenotype in both cell lines.

We reasoned that if *C1QTNF1-AS1* and RSRC2 work together to ensure the correct chromosome alignment, their co-depletion should not aggravate the mitotic phenotype. Accordingly, while the depletion of either *C1QTNF1-AS1* or RSRC2 alone led to the expected delay in mitotic duration, the simultaneous depletion of *C1QTNF1-AS1* and RSRC2 did not have a synergistic effect (Fig. 2 J, K). This data indicates that *C1QTNF1-AS1* acts via the same pathway as RSRC2 to regulate cell division.

### RSRC2 is present in nuclear speckles

RSRC2 (arginine and serine-rich coiled-coil 2) contains an arginine- and serine-rich region, a coiled-coil domain, and a small acidic protein-like (SMAP) domain at its C-terminus (Fig. 3A). Although RSRC2 was identified as an mRNA-interacting protein^34^ and was shown to inhibit proliferation of cancer cell lines^40, 41^, its function is poorly characterised. RSRC2 contains long N-terminal stretches of amino acids that form predicted intrinsically disordered regions (IDRs) (Fig. 3B), which can mediate RNA binding in non-canonical RBPs^14^. Since the serine and arginine-rich (SR) protein class of splicing factors also contain arginine and serine amino acid residues, we hypothesised that RSRC2 is involved in the regulation of splicing. We found RSRC2 to be primarily nuclear, with fluorescence intensity decreasing after its siRNA-mediated depletion (Fig. 3C, D). RSRC2 formed distinct patches in the nuclei resembling nuclear speckles, which are interchromatin granule clusters containing RNAs and splicing factors such as SC35, SON, and SRRM2^42^. To test whether RSRC2 is present within nuclear speckles, we co-stained the cells with antibodies against RSRC2 and SC35, a common nuclear speckles marker^42^. The Manders colocalisation coefficient between the two fluorophores was 0.5347 showing 50% colocalisation of these two channels within the nucleus (Fig. 3E). The colocalisation was reduced in RSRC2-depleted cells (Fig. 3F). Based on its colocalisation within nuclear speckles, the presence of IDRs, and its role as an mRNA-interacting protein^34^, we suggest that RSRC2 could be a novel splicing regulator.

**Figure 3.**
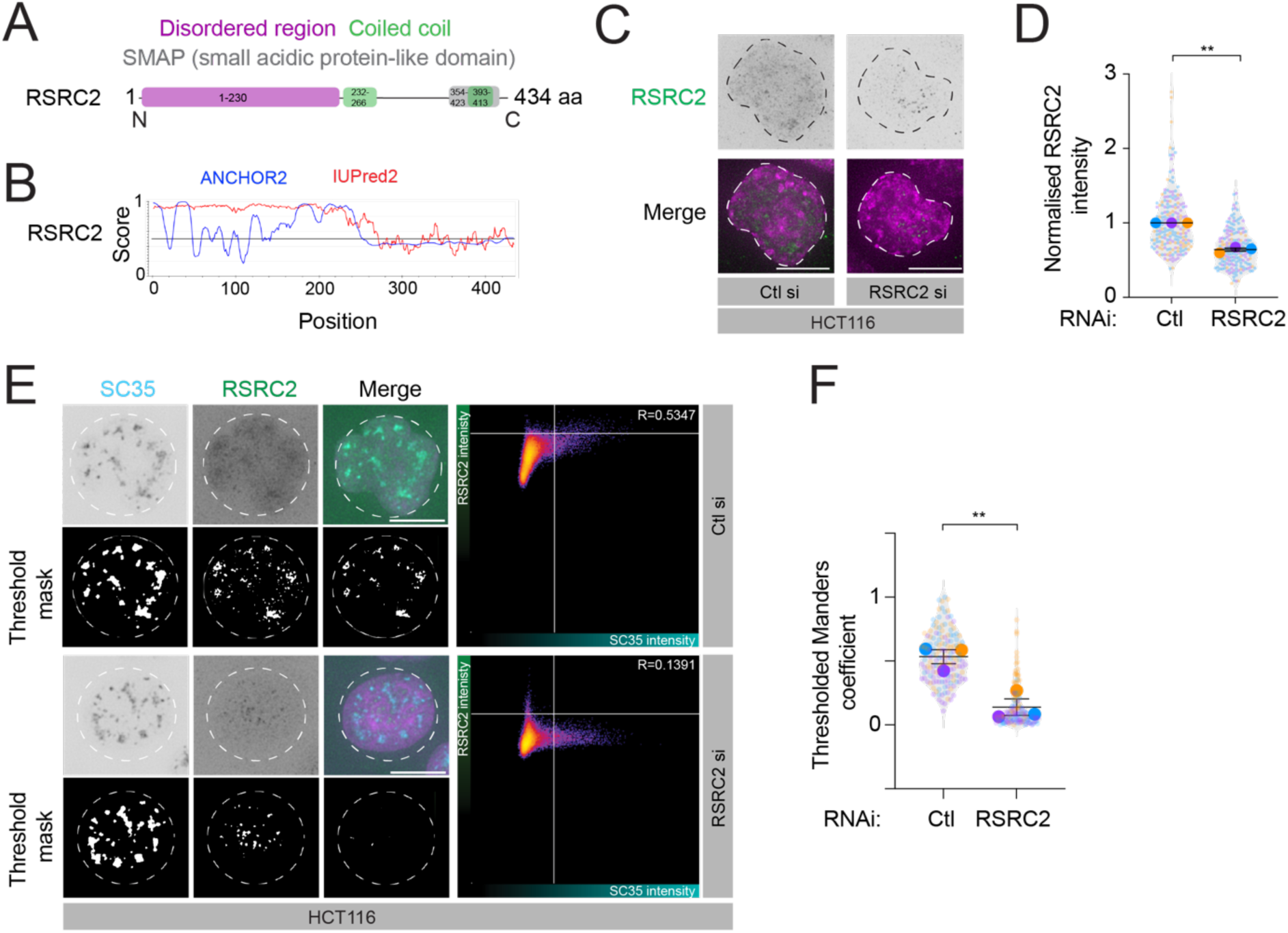
RSRC2, an arginine serine-rich protein, is present in nuclear speckles. (A) Schematic representation of the annotated domain structure of human RSRC2 protein. RSRC2 contains an N-terminal disordered region, two coiled-coil domains and a C-terminal small acidic protein-like (SMAP) domain. (B) Computational disorder prediction for the human RSRC2 protein sequence using IUPred2A (https://iupred2a.elte.hu/) to predict the likelihood of disorder for RSRC2 through the ANCHOR2 or IUPred2 algorithms. Higher disorder scores denote a greater likelihood of the protein region being intrinsically disordered. (C) Representative IF images of interphase HCT116 cells stained with RSRC2 (green) and DNA (Hoescht, magenta) after siRNA-mediated depletion of RSRC2. (D) Quantification of RSRC2 nuclear intensity in HCT116 cells after siRNA-mediated depletion of RSRC2. Results are presented relative to control siRNA (Ctl) treated cells. N = 3 (n_(Ctl si)=_202; n_(RSRC2 si) =_179). (E) Representative images of interphase HCT116 cells stained for nuclear speckles (SC35, cyan) and RSRC2 (green) after siRNA-mediated depletion of RSRC2. DAPI is shown in magenta. Threshold masks used for colocalisation analysis are shown. Correlation plot shows level of correlation between thresholded SC35 and thresholded RSRC2 signal with average Manders correlation coefficient for each condition. (F) Quantitative colocalisation analysis of SC35 and RSRC2 in HCT116 cells following siRNA-mediated depletion of RSRC2 (as shown in panel E). N = 3 (n_(Ctl si)=_154; n_(RSRC2si)=_153). Error bars in all panels are shown as mean ± S.E.M. Scale bar, 5 μm. N = number of cells analysed. The following statistics were applied: one sample t-test (comparing to a hypothetical mean of 1) was used in panel (D); and an unpaired t-test in panel (F).

### RSRC2 protein but not *C1QTNF1-AS1* is a splicing regulator

To investigate whether the *C1QTNF1-AS1*-RSRC2 interaction during cell division involved changes in gene expression levels and alternative splicing, we performed RNA sequencing (RNA-seq) following RNAi and LNA-mediated depletion of RSRC2 and *C1QTNF1-AS1* in HCT116 cells, respectively. We selected differentially expressed genes (DEGs), with adjusted p < 0.01 and logFC > |1|, that consistently changed in the same direction with three LNA gapmers targeting *C1QTNF1-AS1.* This approach was shown to minimise any method-specific off-target effects^7^. RNA-seq analysis revealed that the depletion of *C1QTNF1-AS1* with LNA 1, 2 and 3 gapmers led to 78, 5, and 96 DEGs, respectively (Fig. 4A; Supp. Table 2). The intersection of DEGs with the three LNA gapmers identified no common genes, likely due to the low number of DEGs identified with LNA gapmer 2 (Fig. 4B). Gene ontology (GO) enrichment analysis of RNA-seq datasets after the depletion of *C1QTNF1-AS1* uncovered enrichment in developmental processes, cell differentiation and the G2/M phase of the cell cycle (Supp Fig. 4A; Supp. Table 3). Among the 10 genes identified as potential *C1QTNF1-AS1* targets, *DIAPH2*, *PLK2*, and *PTPRG* are known to have mitotic functions^25, 43^ (Fig. 4C). Further verification of selected targets using qPCR revealed that only DIAPH2 was downregulated following *C1QTNF1-AS1* depletion with two LNAs (Supp. Fig. 4B), which we also confirmed by Western blot (Supp. Fig. 4C, D), highlighting the importance of validating RNA-seq data. In contrast, *DIAPH2* mRNA and protein expression were not affected following RSRC2 knockdown (Supp Fig. 4E-G). Thus, *DIAPH2* is not a shared target between *C1QTNF1-AS1* and RSRC2. Given that we did not observe a major effect on DEGs, we concluded that *C1QTNF1-AS1* does not have major transcriptional roles in HCT116 cells, further supporting our similar observation from HeLa cells^7^. Next, we examined changes in gene expression in cells depleted of RSRC2 compared to control siRNA-transfected cells and identified 259 DEGs in RSRC2-depleted cells (Fig. 4D; Supp. Table 4). GO term annotation in RSRC2-depleted cells identified similar categories, such as cell development and cell differentiation (Supp Fig. 4H; Supp. Table 5). The intersection of DEGs identified in *C1QTNF1-AS1*- and RSRC2-depleted cells did not reveal any shared DEGs (Fig. 4E), indicating that *C1QTNF1-AS1* and RSRC2 do not co-regulate gene expression at the level of mRNA abundance.

**Figure 4.**
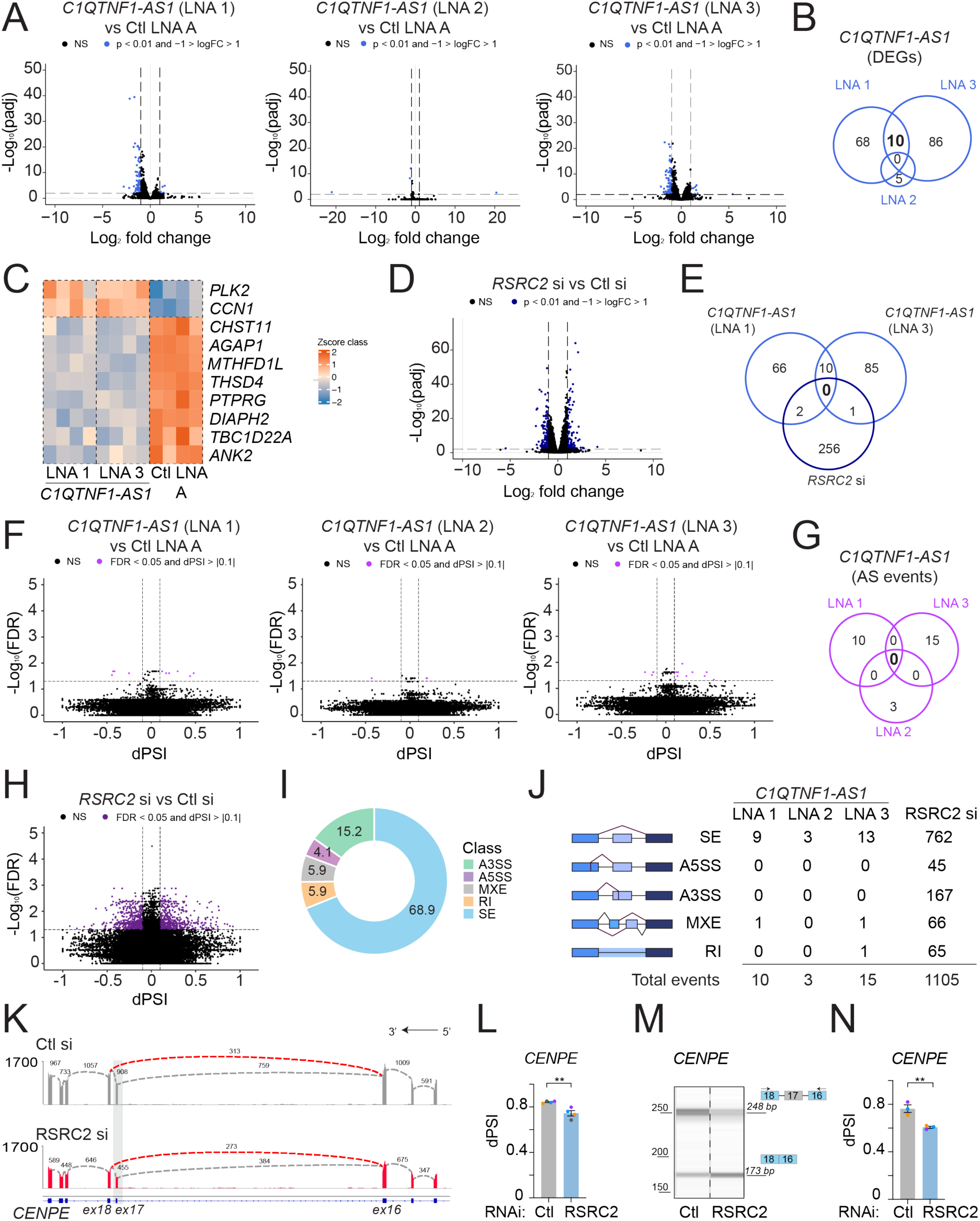
RSRC2, but not *C1QTNF1-AS1*, regulates alternative splicing of genes enriched for organelle and actin cytoskeleton functions. (A) Volcano plot of transcriptional differences induced by LNA-mediated depletion of *C1QTNF1-AS1* in HCT116 cells compared to negative control LNA A (Ctl LNA A). The black horizontal line represents the significance threshold corresponding to a padj of 0.01. Black vertical lines are log_2_-fold change thresholds of > |1|. The genes that do not show significant changes in expression are shown in black. (B) Venn diagram illustrating overlap of the sets of DEGs following depletion of *C1QTNF1-AS1* in HCT116 cells with LNA 1, LNA 2 and LNA 3 targeting *C1QTNF1-AS1.* Only 10 DEGs were in common between LNA 1 and LNA 3 targeting *C1QTNF1-AS1* using the log_2_-fold change threshold of > |1| and FDR of 1%. (C) Heat map of 10 common DEGs from LNA 1- and LNA 3-mediated depletion of *C1QTNF1-AS1* (as shown in panel B). (D) Volcano plot of transcriptional differences induced by siRNA-mediated depletion of RSRC2 in HCT116 cells compared to negative control siRNA. The black horizontal line represents the significance threshold corresponding to a padj of 0.01. Black vertical lines are log_2_-fold change thresholds of > |1|. The genes that do not show significant changes in expression are shown in black. (E) Venn diagram illustrating no overlap between the sets of DEGs identified after siRNA- and LNA-mediated depletion of RSRC2 and *C1QTNF1-AS1* in HCT116 cells. LNA 2 targeting *C1QTNF1-AS1* was omitted in this analysis due to the low number of DEGs. (F) Volcano plot of FDR (log10) versus dPSI of all alternative splicing (AS) events upon *C1QTNF1-AS1* depletion in HCT116 cells. Differential splicing analysis of RNA-seq data was performed with the rMATS (FDR < 0.05 and dPSI > |0.1|). The genes that do not show significant changes in alternative splicing are shown in black. (G) Venn diagram showing no overlap of alternative splicing events following LNA-mediated depletion of *C1QTNF1-AS1* in HCT116 cells. (H) Volcano plot of all alternative splicing events upon RSRC2 silencing in HCT116 cells (as in panel F). (I) Pie chart showing the distribution of types of AS events enriched in RSRC2-depleted HCT116 cells based on rMATS analysis. A3SS – alternative 3′ splice site; A5SS – alternative 5′ splice site; MXE – mutually exclusive exons; RI – retained intron; SE – skipped exon. (J) Table categorising the number of total significant alternative splicing events following *C1QTNF1-AS1*- and RSRC2 depletion in HCT116 cells (FDR < 0.05 and dPSI > |0.1|). (K) Sashimi plot showing skipping of *CENPE* exon 17 (skipped exon is highlighted with a grey rectangle) in RSRC2-depleted HCT116 cells. The tracks were constructed from averages of 4 biological replicates for each condition (RSRC2 si in red; Ctl si in grey). (L) Percentage splicing inclusion (PSI) of *CENPE* exon 17 as calculated from RNA-Seq data of RSRC2-depleted HCT116 cells (as shown in panel K). N = 4. (M) RT-PCR analysis of exon skipping in *CENPE* after RSRC2-mediated depletion compared to control siRNA. PCR products were amplified using primers against *CENPE* long and short isoforms, with and without exon 17, and were visualised using capillary gel electrophoresis. (N) Quantification of PSI for *CENPE* exon 17 based on capillary gel electrophoresis (shown in panel M) following RSRC2 depletion. N = 3. Error bars in panels (L) and (N) are shown as mean ± S.E.M. Statistics were applied using an unpaired t-test in panels (L) and (N).

Since RSRC2 was present in the nuclear speckles, we considered that RSRC2 and *C1QTNF1-AS1* could be involved in gene regulation at the level of splicing and therefore co-regulate a common set of transcripts with functions in mitosis. Thus, we performed differential splicing analysis using rMATS^44^ and searched for mis-regulated splicing events in *C1QTNF1-AS1*- and RSRC2-depleted cells. Our analyses identified 10, 3, and 15 alternative splicing events (FDR < 0.05, |ΔPSI| > 0.1, where PSI is the Percentage Splicing Inclusion) following *C1QTNF1-AS1* depletion using LNA 1, 2, and 3 gapmers, respectively (Fig. 4F). Intersection of alternative splicing events upon *C1QTNF1-AS1* depletion with three LNA gapmers revealed no shared mis-regulated splicing variations (Fig. 4G; Supp. Table 6). Given that *C1QTNF1-AS1* is a low-abundant lncRNA and the mis-splicing events upon its loss were minor, we conclude that *C1QTNF1-AS1’s* role in cell division does not occur through alternative splicing. We then turned our attention to RSRC2 and examined its role in splicing regulation.

Using rMATS analysis to compare splice variations from RSRC2-depleted cells, we identified 1,105 RSRC2-dependent alternative splicing events (Fig. 4H; Supp Table 7), with exon skipping and alternative 3′ splice sites being the most frequently occurring alternative splice events (Fig. 4I). GO analysis of 876 differentially spliced genes indicated that RSRC2-regulated transcripts are involved in organelle and cytoskeletal organisation as some of the top enriched pathways (Supp Fig. 4I; Supp. Table 8), which could explain the mitotic defects observed in RSRC2-depleted cells. Furthermore, analysis of all the significant pathways associated with the RSRC2 mis-spliced genes indicated that some transcripts are linked to the regulation of the cell cycle, microtubule polymerisation and centrosome (Supp Fig. 4J). By comparing the number of differential alternative splicing events between RSRC2- and *C1QTNF1-AS1*-depleted cells, we further confirmed that the loss of *C1QTNF1-AS1* did not lead to splicing changes, in contrast to RSRC2 depletion (Fig. 4J).

As aberrant splicing of *CENPE* mRNA (centrosome-associated protein E), a kinesin-like motor protein, has been reported to lead to chromosome congression defects^45^, we analysed its splicing upon RSRC2 depletion. We found *CENPE* among the alternatively spliced candidates in RSRC2-depleted cells, cross-validating our splicing analysis. As expected, the sashimi plot confirmed an exon-skipping event in exon 17 of *CENPE* following RSRC2 depletion (Fig 4. K, L; dPSI = −0.1). Additionally, RT-PCR analysis validated the *CENPE* exon splicing by capillary gel electrophoresis (Fig. 4 M, N). Thus, RSRC2-dependent regulation of *CENPE* pre-mRNA splicing could explain the origin of chromosome congression defects observed in RSRC2-depleted cells. Together, these data indicate that RSRC2 is a novel splicing regulator modulating the splicing of numerous genes involved in cell cycle progression independently of *C1QTNF1-AS1*.

### RSRC2 interacts with a set of proteins involved in splicing and cell division

To gain insights into the molecular mechanisms of how RSRC2 regulates alternative splicing, we conducted an Immunoprecipitation coupled with Mass-Spectrometry (IP-MS) to identify either RNA-mediated protein-protein or direct protein-protein interactions of RSRC2. As shown for other RBPs using similar approach^46^, RSRC2 was efficiently and specifically immunoprecipitated from whole-cell extracts from HCT116 cells, compared to a matched IgG control antibody, in the presence and absence of RNase A/T1 (Suppl. Fig. 5A). We confirmed that RNase A/T1 was effective in degrading the RNA, compared to whole-cell extracts treated without RNase A/T1 (Suppl. Fig. 5B). The analysis of the RSRC2 interactome without RNase A/T1 identified 123 proteins (Fig. 5A; Supp. Table 9) that were significantly enriched in the RSRC2 IP compared to IgG controls. GO analysis of the RSRC2 interactors demonstrated a distinct enrichment of proteins associated with splicing, centrosome, ribosome biogenesis and cell adhesion (Fig. 5B). Indeed, some of the RSRC2-protein interactors included proteins involved in the regulation of splicing and nuclear speckle assembly such as SRRM2 and SON^42, 47^. Among the top RSRC2 protein interactors, we also found centrosome proteins, such as pericentrin (PCNT) and cyclin-dependent kinase 5 regulatory subunit associated protein 2 (CDK5RAP2/CEP215). PCNT and CDK5RAP2 are key pericentriolar material (PCM) proteins that, together with the centrioles, form a centrosome, a major microtubule-organising centre that contributes to mitotic fidelity^48, 49^. The interaction of RSRC2 with splicing and centrosome proteins among its key interactors suggests that RSRC2 could assemble multiple ribonucleoprotein complexes to perform various functions across different subcellular locations.

**Figure 5.**
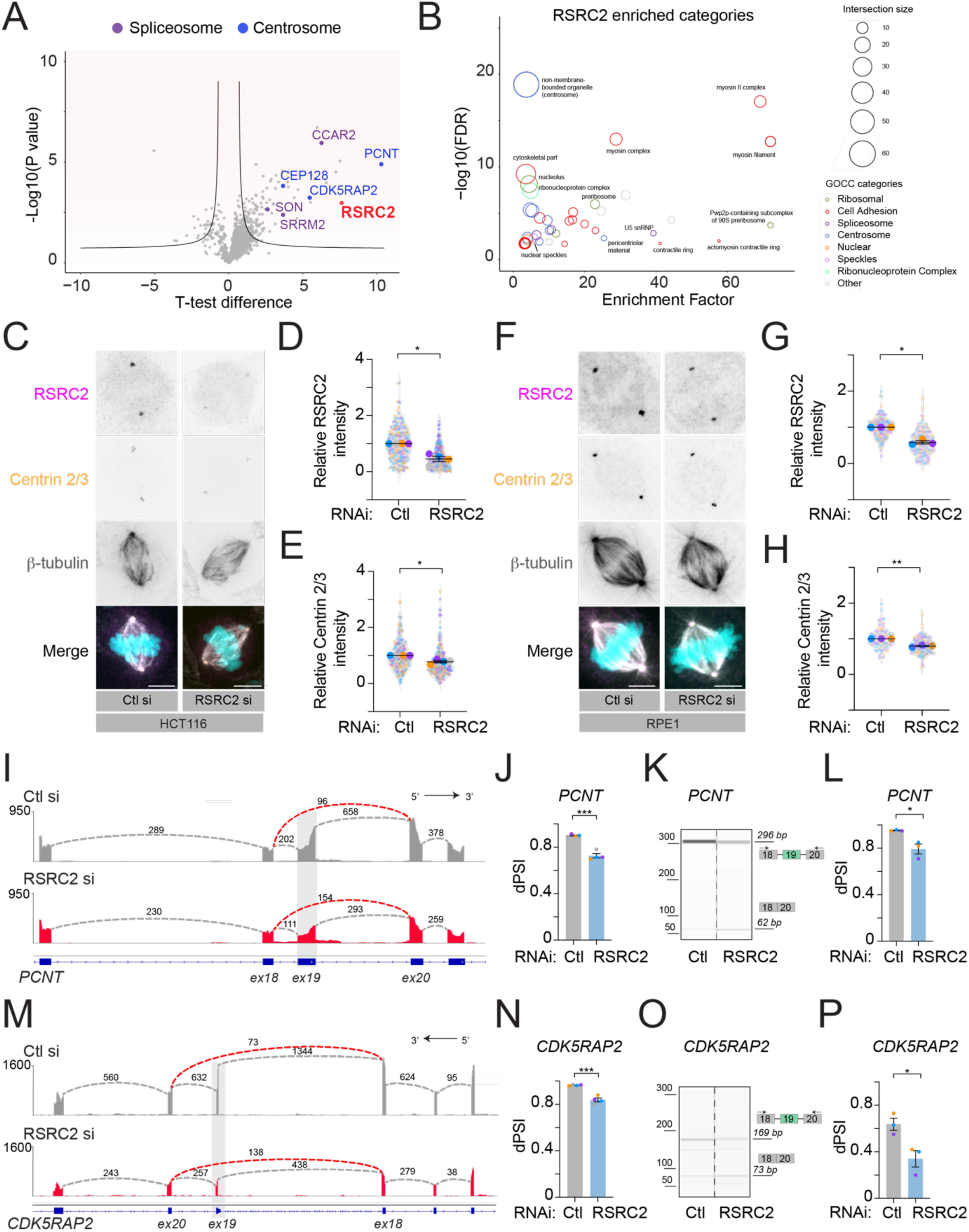
Identification of the RSRC2 interactome uncovered the RSRC2 protein-protein interaction networks, which include many proteins that mediate cell division and splicing. (A) Volcano plot of protein-protein interactions for RSRC2 comparing enriched proteins of the RSRC2 IP versus IgG IP in HCT116 cells without RNase A/T1 treatment. Curved lines mark the significance boundary (FDR = 1%, at least two peptides detected in each biological replicate). Each dot represents a protein. Selected splicing and centrosome proteins are indicated as one of the top RSRC2 interactors. N = 4. Statistically significant differences were detected using a two-tailed, two-sample t-test with permutation-based FDR. (A) Fisher’s exact test analysis of known protein categories that are over-represented among the RSRC2 interacting proteins (Benjamini–Hochberg FDR < 0.05). Each circle represents an enriched category from the Gene Ontology Cellular Compartments (GOCC) database, with the circle size representing the number of shared proteins. (B) Representative images of mitotic HCT116 cells stained for RSRC2 (magenta), centrosomes (Cen2/3, orange), microtubules (β-tubulin, grey) and DNA (Hoescht, cyan) after siRNA-mediated depletion of RSRC2. (C) Quantification of centrosomal RSRC2 intensity in HCT116 cells from maximum intensity projections obtained from panel (C). Results are presented relative to control siRNA (Ctl) treated cells. N = 4 (n_(Ctl si)=_221; n_(RSRC2 si))=_226). (D) Quantification of centrosomal Cen2/3 intensity in HCT116 cells from maximum intensity projections obtained from panel (C). Results are presented relative to control siRNA (Ctl) treated cells. N = 4 (n_(Ctl si)=_220; n_(RSRC2 si))=_231). (E) Representative images of mitotic RPE1 cells stained as in panel (C) after siRNA-mediated depletion of RSRC2. (F) Quantification of centrosomal RSRC2 intensity in RPE1 cells from maximum intensity projections obtained from panel (F). Results are presented relative to control siRNA (Ctl) treated cells. N = 3 (n_(Ctl si)=_133; n_(RSRC2 si))=_124). (G) Quantification of centrosomal Cen2/3 intensity in RPE1 cells from maximum intensity projections obtained from panel (F). Results are presented relative to control siRNA (Ctl) treated cells. N = 3 (n_(Ctl si)=_133; n_(RSRC2 si))=_124). (H) Sashimi plot showing skipping of *PCNT* exon 19 (skipped exon is highlighted with a grey rectangle) in RSRC2-depleted HCT116 cells. The tracks were constructed from averages of 4 biological replicates for each condition (RSRC2 si in red; Ctl si in grey). (I) Percentage splicing inclusion (PSI) of *PCNT* exon 19 was calculated from RNA-seq data of RSRC2-depleted HCT116 cells (as shown in panel I) and the difference was −0.17421 (dPSI). N = 4. (J) RT-PCR analysis of exon skipping in *PCNT* after RSRC2-mediated depletion compared to control siRNA. PCR products were amplified using primers against *PCNT* and were visualised using capillary gel electrophoresis. (K) Quantification of dPSI for *PCNT* exon 19 based on capillary gel electrophoresis (shown in panel K) following RSRC2 depletion. N = 3. (L) Sashimi plot showing skipping of *CDK5RAP2* exon 19 (skipped exon is highlighted with a grey rectangle) in RSRC2-depleted HCT116 cells. The tracks were constructed from averages of 4 biological replicates for each condition (RSRC2 si in red; Ctl si ingrey). (M) Percentage splicing inclusion of *CDK5RAP2* exon 19 was calculated from RNA-seq data of RSRC2-depleted cells (as shown in panel M) and was −0.12747 (dPSI). N = 4. (N) RT-PCR analysis of exon skipping in *CDK5RAP2* after RSRC2-mediated depletion compared to control siRNA. PCR products were amplified using primers against *CDK5RAP2* and were visualised using capillary gel electrophoresis. (O) Quantification of dPSI for *CDK5RAP2* exon 19 based on capillary gel electrophoresis (shown in panel O) following RSRC2 depletion. N = 3. Error bars in all panels are shown as mean ± S.E.M. Scale bar, 5 μm. N = number of cells analysed. For panels (D), (E), (G) and (H), a one-sample *t*-test (using a hypothetical mean of 1) was used. An unpaired t-test was used in panel (J), (L), (N) and (P).

To determine if the association of RSRC2 with splicing and centrosome proteins was RNA-mediated, we repeated the IP-MS in the presence of RNase A/T1. The interaction profile of RSRC2 was similar to that with no RNase A/T1 treatment (Supp. Fig. 5C, Supp. Table 9), demonstrating that the majority of RSRC2 protein-protein interactions are RNA-independent. Furthermore, GO analysis of RSRC2 interactors following RNase A/T1 treatment showed similar enrichment in pathways associated with RSRC2 in presence of RNA (Supp. Fig. 5D) suggesting that the majority of RSRC2 protein-protein interactions are RNA-independent. Combined, these data and our findings that loss of RSRC2 leads to mitotic defects suggest that RSRC2 is a multifunctional protein regulating centrosome biogenesis and splicing. To validate this, we performed IF in control- and RSRC2-depleted cells by staining the cells with RSRC2 and centrin (Cen 2/3), a centriole marker. We found RSRC2 to be present on the centrosomes of mitotic cells, while RSRC2-centrin colocalisation was reduced following RSRC2 depletion in HCT116 (Fig. 5C, D) and RPE1 cells (Fig. 5F, G). We also observed a significant reduction of the centrin levels in HCT116 and RPE1 cells upon RSRC2 loss (Fig. 5H, E), indicating that RSRC2 may regulate centriole assembly during mitosis.

In view of our rMATS data indicating that several RSRC2 alternative spliced genes were linked to pathways associated with centrosomes, we considered that *PCNT* and *CDK5RAP2* could also be potential RSRC2 splicing targets. Our splicing analysis showed exon skipping for *PCNT* in RSRC2-depleted cells, which can induce a premature stop codon (Fig. 5I, J). Capillary gel electrophoresis verified that *PCNT* exon 19 is skipped in RSRC2-depleted cells (Fig. 5K, L). Additionally, RSRC2 knockdown led to exon skipping of *CDK5RAP2* exon 19 (Fig. 5M, N), which was further verified using the capillary gel electrophoresis in a separate set of experiments (Fig. 5O, P). Thus, RSRC2 has a role in the splicing of a specific group of genes associated with mitosis.

To assess the contribution of RSRC2 splicing targets to the chromosome congression defects observed in RSRC2-depleted cells, we examined CREST and α-tubulin staining upon PCNT knockdown. Consistent with the previous report^50^, PCNT-depleted cells showed an increase in chromosome congression defects compared to control siRNA-treated cells (Supp. Fig. 5 E, F; Fig. 6 G). Therefore, the resulting splicing defects in *PCNT* —along with other mitotic targets of RSRC2 like *CENPE* (Fig. 4K-N)—are likely the cause of the chromosome congression defects observed in RSRC2-depleted cells. Together, these results indicate that RSRC2 is a novel splicing factor required for maintaining the homeostasis of mRNAs that code for centrosome proteins.

**Figure 6.**
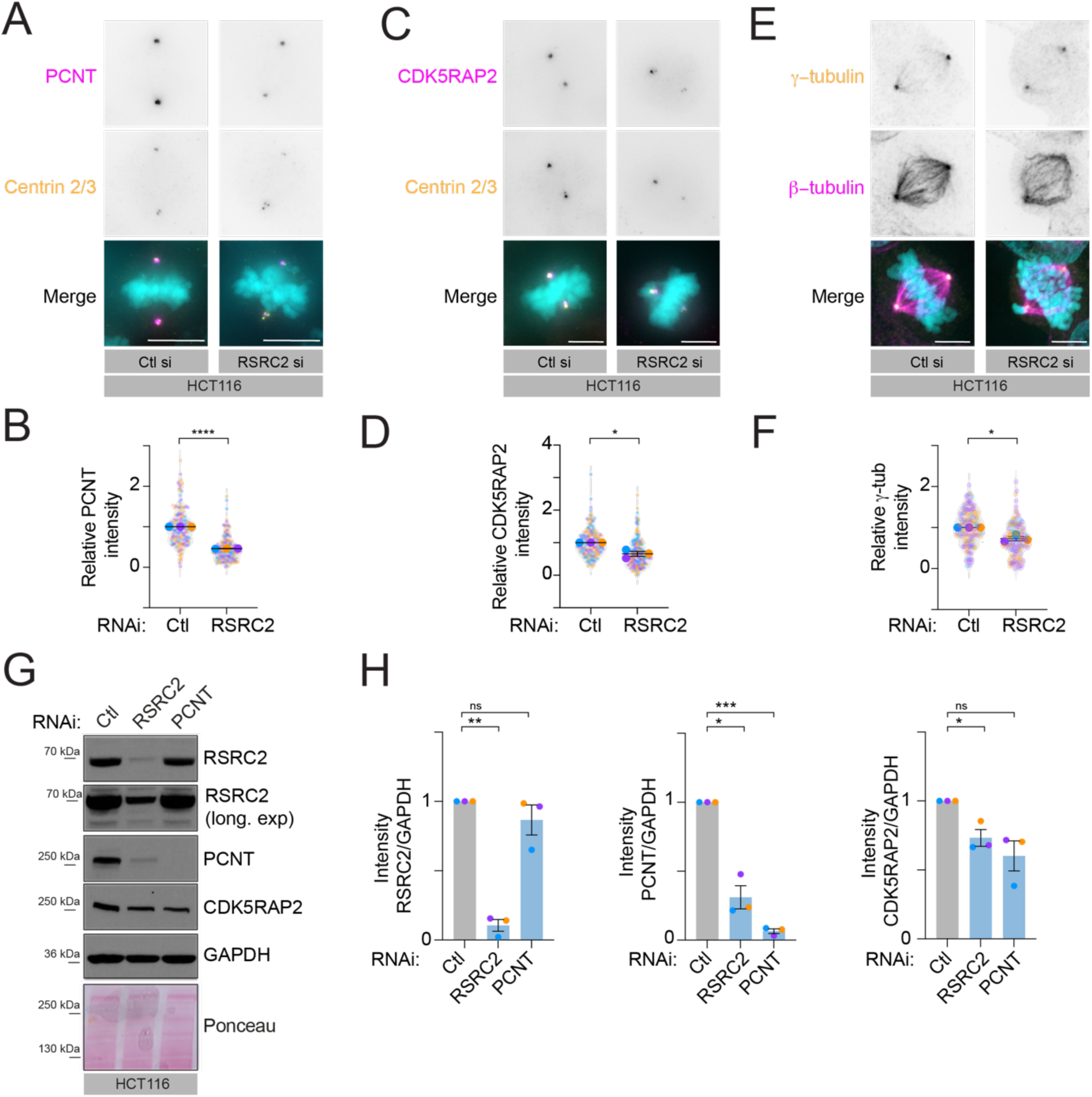
Depletion of RSRC2 reduces the levels of PCNT, CDK5RAP2 and γ-tubulin at centrosomes of mitotic cells. (A) Representative images of mitotic HCT116 cells stained for PCM (PCNT, magenta), centrioles (Cen2/3, orange) and DNA (Hoescht, cyan) after siRNA-mediated depletion of RSRC2. Centrosome contains a pair of centrioles and surrounding PCM matrix (PCNT, CDK5RAP2). (B) Quantification of PCNT signal intensity around the centrosome of mitotic HCT116 cells from the maximum intensity projections obtained from panel (A). Results are presented relative to control siRNA (Ctl) treated cells. N = 3 (n_(Ctl si)=_169; n_(RSRC2 si))=_161). (C) Representative images of mitotic HCT116 cells stained for PCM (CDK5RAP2, magenta), centrioles (Cen2/3, orange) and DNA (Hoescht, cyan) after siRNA-mediated depletion of RSRC2. (D) Quantification of CDK5RAP2 signal intensity around the centrosome of mitotic HCT116 cells from the maximum intensity projections obtained from panel (C). Results are presented relative to control siRNA (Ctl) treated cells. N = 3 (n_(Ctl si)=_176; n_(RSRC2 si))=_180). (E) Representative images of mitotic HCT116 cells stained for centrosomes (γ-tubulin, orange), microtubules (β-tubulin, magenta) and DNA (Hoescht, cyan) after siRNA-mediated depletion of RSRC2. (F) Quantification of γ-tubulin signal intensity around the centrosome of mitotic HCT116 from maximum intensity projections obtained from panel (E). Results are presented relative to control siRNA (Ctl) treated cells. N = 3 (n_(Ctl si)=_143; n_(RSRC2 si))=_137). (G) Representative western blot of HCT116 cells probed with RSRC2, PCNT, CDK5RAP2 and GAPDH antibodies after the RSRC2 and PCNT knockdown. GAPDH and Ponceau staining were used as loading controls. (H) Densitometric analysis of RSRC2, PCNT and CDK5RAP2 levels from panel (G) relative to control siRNA (Ctl). N = 3. Error bars in all panels are shown as mean ± S.E.M. Scale bar, 5 μm. N = number of cells analysed. For panels (B), (D) and (F), a one-sample *t*-test (comparing to a hypothetical mean of 1) was used. An unpaired t-test with Welch’s correction was used in panel (H).

### RSRC2 is required for the recruitment of PCNT, CDK5RAP2 and γ-tubulin on mitotic centrosomes

Having demonstrated that RSRC2 is required for efficient pre-mRNA splicing of genes involved in mitosis, we evaluated the consequences of *PCNT* and *CDK5RAP2* mis-splicing in RSRC2-depleted cells. RSRC2-depleted cells exhibited 45% and 65% reduction of PCNT (Fig. 6 A, B) and CDK5RAP2 (Fig. 6 C, D) localisation at the mitotic centrosomes respectively, compared to control siRNA-treated cells. Since PCNT is required for tethering the γ-tubulin ring complex (γ-TuRC) and is essential for microtubule nucleation from centrosomes in mitosis^51, 52^, we examined if RSRC2 loss affects centrosome-localised γ-tubulin. As predicted, the loss of RSRC2 led to a 27% reduction of γ-tubulin at centrosomes in mitotic cells (Fig. 6 E, F). Consistent with the reduced PCNT and CDK5RAP2 levels on mitotic centrosomes, the Western blot confirmed that RSRC2 is required for maintaining the protein levels of PCNT and CDK5RAP2 (Fig. 6 G, H). We also observed diminished protein levels of CDK5RAP2 in PCNT-depleted cells. This is in line with the observation that CDK5RAP2 interacts with PCNT^53^ and that loss of PCNT leads to a reduction in centrosomal CDK5RAP2^54, 55^. Collectively, these results indicate that RSRC2 is required for the distribution of PCNT, CDK5RAP2, and γ-tubulin at mitotic centrosomes as well as their protein abundance. Since our splicing analysis showed that the loss of RSRC2 causes exon skipping in *PCNT* and *CDK5RAP2*, we propose that this reduction in protein levels of PCNT and CDK5RAP2 diminishes their recruitment to centrosomes, ultimately affecting their mitotic function.

### *C1QTNF1-AS1* depletion alters RSRC2 localisation on centrosomes

Recently, the biological significance of RNA-protein interactions has shifted from a protein-focused to a more RNA-centric perspective where RNA can control protein function by affecting protein conformation, intracellular localisation, interaction and enzymatic activity^14, 56–60^. To gain insights into the function of *C1QTNF1-AS1* and RSRC2 interaction during cell division, we examined whether *C1QTNF1-AS1* might affect RSRC2 levels, its interactome, and localisation. Depletion of *C1QTNF1-AS1* did not affect *RSRC2* mRNA levels (Suppl. Fig. 6A), total protein amounts (Suppl. Fig. 6B, C), nor its localisation in the nucleus (Suppl. Fig. 6D, E). Similarly, *C1QTNF1-AS1* expression did not change upon RSRC2 depletion in HCT116 and RPE1 cell lines (Suppl. Fig. 6F, G). Considering that RSRC2’s splicing function was independent of *C1QTNF1-AS1* (Fig. 4) and *C1QTNF1-AS1* did not impact RSRC2 total protein levels or its nuclear localisation, we asked whether *C1QTNF1-AS1* might act as a regulator of RSRC2 mitotic localisation. To test this, we conducted the IF for RSRC2 and centrin in *C1QTNF1-AS1*-depleted cells and found a reduction to 71% *(C1QTNF1-AS1* LNA 1) and 78% reduction in RSRC2 localisation (*C1QTNF1-AS1* LNA 3) on mitotic centrosomes compared to Ctl A (Fig. 7 A, B). Centrin levels did not change upon *C1QTNF1-AS1* knockdown (Fig. 7 C), indicating that *C1QTNF1-AS1* is not involved in centriole biogenesis. Furthermore, loss of *C1QTNF1-AS1* did not affect the protein levels of PCNT and CDK5RAP2 (Supp. Fig. 7 A, B), nor their centrosome localisation on the mitotic spindle (Supp. Fig. 7 C-F). This suggests that *C1QTNF1-AS1* is not required for the distribution of PCNT and CDK5RAP2 at mitotic centrosomes, in contrast to RSRC2.

**Figure 7.**
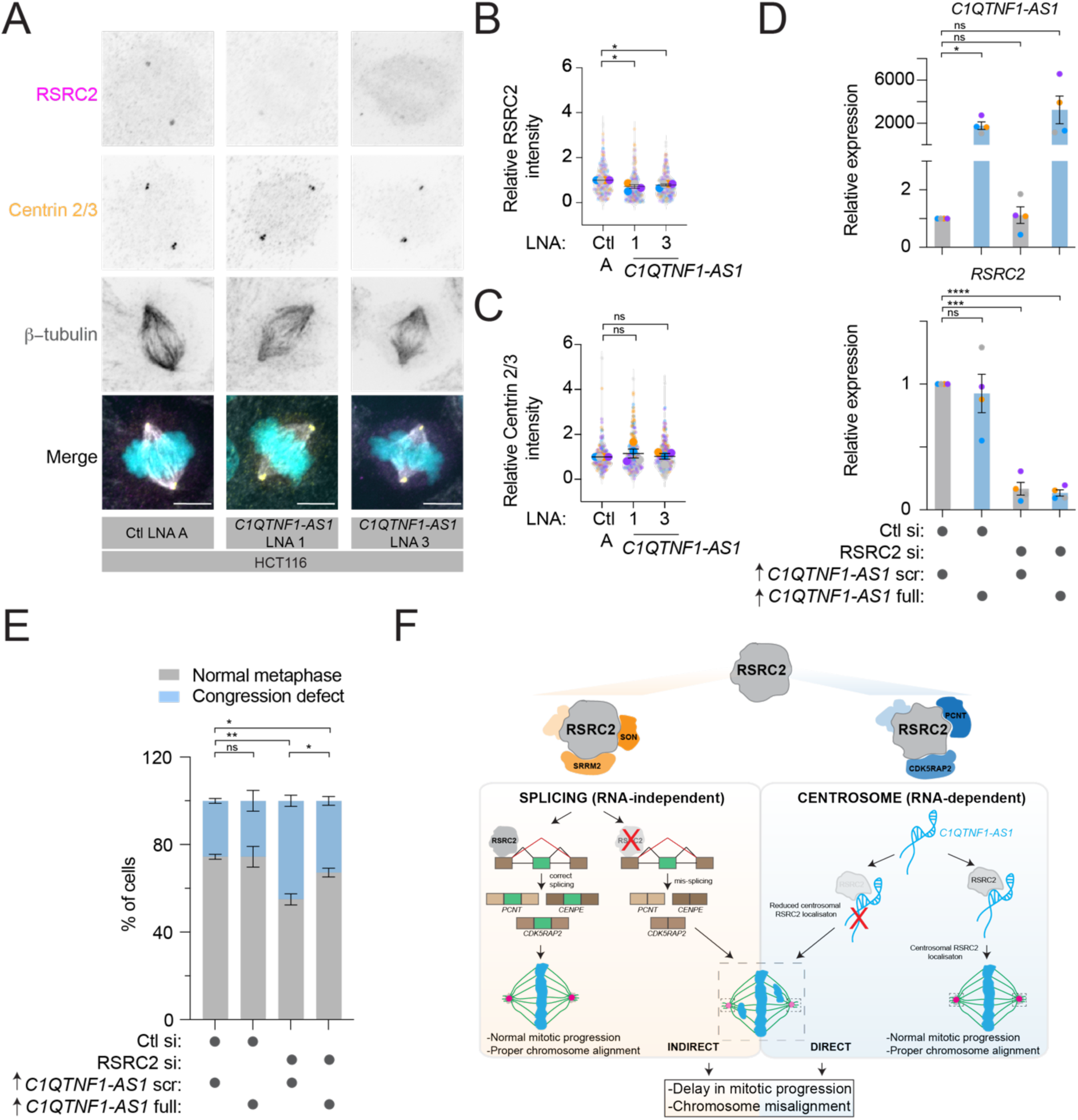
*C1QTNF1-AS1* promotes RSRC2 recruitment at the mitotic centrosomes and restores chromosome congression defects in RSRC2-depleted cells. (A) Representative images of mitotic HCT116 cells stained for RSRC2 (magenta), centrosomes (Cen2/3, orange), microtubules (β-tubulin, grey) and DNA (Hoescht, cyan) after LNA-mediated depletion of *C1QTNF1-AS1*. (B) Quantification of RSRC2 signal intensity around the centrosome of mitotic HCT116 cells from maximum intensity projections obtained from panel (A). Results are presented relative to control LNA A (Ctl A) treated cells. N = 4 (n_(Ctl A)=_204; n_(*C1QTNF1-AS1* LNA1)=_245; n_(*C1QTNF1-AS1* LNA5)=_221). (C) Quantification of Cen2/3 signal intensity around the centrosome of mitotic HCT116 cells from maximum intensity projections obtained from panel (A). Results are presented relative to control LNA A (Ctl A) treated cells. N = 4 (n_(Ctl A)=_217; n_(*C1QTNF1-AS1* LNA1)=_269; n_(*C1QTNF1-AS1* LNA5)=_252). (D) Expression levels of *C1QTNF1-AS1* and RSRC2 in HCT116 cells following depletion of RSRC2 and overexpression of either scrambled (Scr) or full sequences of *C1QTNF1-AS1*, as measured by qPCR. Results are presented relative to negative control siRNA (Ctl si) and *C1QTNF1-AS1* (Scr). N = 4. (E) Characterisation of mitotic phenotypes in HCT116 cells following depletion of RSRC2 and overexpression of either scrambled (Scr) or full sequences of *C1QTNF1-AS1* in HCT116 cells. The percentage of mitotic cells with normal metaphase plate and congression defect was determined using the same antibodies as in Fig. 2A. N = 3 (n_(Ctl si+ *C1QTNF1-AS1 scr*)=_180; n_(Ctl si+ *C1QTNF1-AS1 full*)=_180; n_(RSRC2 si+ *C1QTNF1-AS1 scr*)=_180; n_(RSRC2 si+ *C1QTNF1-AS1 full*)=_180). (F) Current working model of how RSRC2’s dual function contributes to mitotic fidelity. RSRC2 is a multifunctional RBP whose loss leads to mitotic delay and chromosome congression defects. The RSRC2-dependent mitotic phenotype is due to its direct function in mitosis (right panel) by interacting with centrosome proteins such as PCNT and CDK5RAP2, and an indirect role via splicing of pre-mRNAs encoding mitotic regulators (left panel). *C1QTNF1-AS1* contributes to the direct function of RSRC2 by facilitating its recruitment at the mitotic centrosomes. Error bars in all panels are shown as mean ± S.E.M. Scale bar, 5 μm. N = number of cells analysed. The following statistics were applied: one sample t-test (using a hypothetical mean of 1) in panels (B) and (C); and an unpaired t-test with Welch’s correction in (D) and (E).

As lncRNAs are known to regulate protein activities^61^, we next explored whether *C1QTNF1-AS1* may be necessary for maintaining the RSRC2 interactome. First, we performed the IP-MS for RSRC2 in control- and *C1QTNF1-AS1*-depleted cells and found no significant changes among top interactors of RSRC2 (Supp. Fig. 6H; Supp. Table 10). Indeed, PCNT and CDK5RAP2 were still bound to RSRC2 even in the absence of *C1QTNF1-AS1*. Combined with RSRC2 IP-MS with and without RNase A/T1, we show that the majority of RSRC2 protein-protein interactions are RNA-independent (Supp. Fig. 5C). Given that serine/arginine-rich proteins are highly phosphorylated on serine residues, we reasoned that *C1QTNF1-AS1* could influence the phosphorylation of serine (Ser) resides in RSRC2, thereby impacting its localisation and activity, as shown for other splicing factors^62, 63^. We searched for RSRC2 phospho (STY) peptides in control- and *C1QTNF1-AS1*-depleted cells, which revealed that RSRC2 has one threonine (T) and two serine (S) residues that are phosphorylated. However, their phosphorylation status did not change upon *C1QTNF1-AS1* loss (Supp. Fig. 6I; Supp. Table 10). These observations indicate that *C1QTNF1-AS1* does not alter the RSRC2 protein interactome or its phosphorylation status.

To ascertain whether *C1QTNF1-AS1*-dependent localisation of RSRC2 on centrosomes is necessary for faithful mitosis, we performed IF in RSRC2-depleted cells upon overexpression of *C1QTNF1-AS1*, using either its full or scrambled sequences (Fig. 7D). Overexpression of the mature *C1QTNF1-AS1* transcript but not its scrambled sequence variant rescued the chromosome congression defects in RSRC2-depleted cells (Fig. 7 E). As the siRNA-mediated depletion of RSRC2 was not complete as shown by Western blot (Fig. 1H, 6G) and IF (Fig. 3D), we hypothesise that the residual RSRC2 protein in RSRC2-depleted cells can interact with the overexpressed *C1QTNF1-AS1* and rescue the mitotic phenotypes. Together, these results show that the primary function of *C1QTNF1-AS1* is to regulate RSRC2 localisation on centrosomes to ensure error-free mitosis.

## Discussion

In this study, we discovered RSRC2 as a novel regulator of cell division that acts as a dual-functional protein. The primary function of RSRC2 is to act as a splicing regulator for a set of mitotic genes, while its other role involves recruitment to the mitotic centrosomes, which requires interaction with lncRNA *C1QTNF1-AS1*.

First, we show that the mechanism by which *C1QTNF1-AS1* controls cell division involves its interaction with the RSRC2 RNA-binding protein. Our findings demonstrate that loss of both *C1QTNF1-AS1* and RSRC2 leads to chromosome congression defects and mitotic delay. We further reveal that RSRC2, but not *C1QTNF1-AS1*, is a novel splicing regulator of genes enriched in cell cycle functions. Interestingly, besides splicing interactors, several centrosome proteins, such as PCNT and CDK5RAP2, can interact with RSRC2. While numerous proteins, including splicing regulators^28^, have been shown to moonlight— suggesting they perform multiple functions in different cellular locations^64^—the regulation of location and timing of their activity remains unclear. Our data indicate that RSRC2 has a dual mitotic function, an indirect function required for splicing of mitotic regulators, and a direct one through its localisation to the mitotic centrosomes and interaction with centrosome proteins (Fig. 7F).

Second, we found that the dual function of RSRC2 can be RNA-mediated. We provide evidence that the splicing-related function of RSRC2 is independent of *C1QTNF1-AS1*, whereas *C1QTNF1-AS1* mediates RSRC2 recruitment on the mitotic centrosomes to ensure error-free mitosis. This is supported by the double knockdown experiment, as the loss of *C1QTNF1-AS1* and RSRC2 did not show additive effects. Although overexpression of *C1QTNF1-AS1* in RSRC2-depleted cells rescued chromosome congression defects, indicating that RSRC2 may regulate *C1QTNF1-AS1*, we cannot exclude alternative scenarios caused by incomplete siRNA-mediated knockdown of RSRC2. It is possible that the remaining RSRC2 protein in RSRC2-depleted cells binds to *C1QTNF1-AS1* and restores the mitotic phenotype. As *C1QTNF1-AS1* is not present on the centrosomes, this raises the question of how *C1QTNF1-AS1* contributes to RSRC2 localisation? The low copy number of *C1QTNF1-AS1* makes it challenging to study. Thus, it is possible that our imaging techniques did not reach single-molecule sensitivity and consequently failed to capture *C1QTNF1-AS1* on the mitotic spindle. Methods such as 3D super-resolution microscopy could help overcome this limitation, similar to what has been demonstrated with the lncRNA *SLERT*^65^. Given that short RNA motifs and/or structural domains in lncRNAs drive several lncRNA-protein interactions ^31^, one might expect that similar functional elements are present in *C1QTNF1-AS1* that influence RSRC2 localisation. Future experiments using methods such as *in vivo* SHAPE-Map^66^ and/or COMRADES^67^, will shed light on whether RNA structure or short elements within *C1QTNF1-AS1* contribute to the *C1QTNF1-AS1*-RSRC2 interaction to facilitate error-free mitosis.

Our experiments also highlight that RSRC2 is a multifunctional RBP, as shown recently for ∼100 endogenous RBPs^46^. RSRC2 localises to nuclear speckles and interacts with splicing factors, including SON and SRRM2, which are essential for nuclear speckle formation^42^. Additionally, the loss of RSRC2 affects the mRNA splicing of genes enriched with mitotic functions, such as *CENPE, PCNT* and *CDK5RAP2*. Although we cannot distinguish whether the mis-splicing of *PCNT* and *CDK5RAP2* or RSRC2’s binding to these proteins causes chromosome congression defects in RSRC2-depleted cells, we favour the explanation that the mis-splicing of mitotic genes primarily contributes to the observed defects in these cells. In support of our model, the splicing of *PCNT* pre-mRNA has been shown to be dependent on SON^17, 18^, a protein we identified as part of the RSRC2 interactome. Thus, we propose that the interaction between RSRC2 and SON could regulate the splicing of a specific group of genes associated with mitosis. Despite being listed in the RBP2GO database of RBPs^34^, RSRC2 lacks a well-defined RNA-binding domain and thus belongs to a large number of non-canonical RBPs^14^. Indeed, RSRC2 is classified as non-canonical RBPs in the RBP catalogue known as “RBPWorld”^68^. Follow-up work mapping the RNA-binding landscape of RSRC2 will further elucidate its role as a multifunctional RBP.

We also show that RSRC2 has a direct role in cell division through three lines of evidence: i) RSRC2 localises to centrosomes, ii) RSRC2 physically interacts with centrosome proteins such as PCNT and CDK5RAP2, and iii) RSRC2 loss leads to mitotic delay and chromosome congression defects. Thus, our study adds to the growing literature that moonlighting in mitosis is a shared characteristic of many splicing regulators^28^, including, now also, RSRC2. Notably, the loss of RSRC2 leads to a reduction of PCNT, CDK5RAP2 and γ-tubulin at mitotic centrosomes, suggesting that RSRC2 loss is mimicking the lack of centrosomes. Due to the absence of centrosomes, which facilitate mitotic fidelity^48^, it is possible that RSRC2-depleted cells exhibit defects in microtubule organisation and dynamics, leading to chromosome misalignment. Finally, the dual role of RSRC2 in cell division is further supported by the RSRC2 domain structures. In addition to IDRs that are known to interact with RNA, RSRC2 also contains coiled-coil domains that are present in many centrosome proteins, including PCNT and CDK5RAP2. We reasoned that RSRC2 could contribute to the overall PCM architecture of the centrosomes through these coiled-coil domains^49^.

In summary, we have identified RSRC2 as a *C1QTNF1-AS1* interactor and a novel regulator of mitosis. Furthermore, our study on the *C1QTNF1-AS1* and RSRC2 interaction highlights the importance of lncRNAs in regulating the mitotic, but not the splicing function of RSRC2. We believe our research will open new avenues in understanding how (lnc)RNA-protein interactions promote error-free mitosis.

## Methods

### Cell culture

Human normal retinal pigment epithelial hTERT-RPE1 (ATCC) and hTERT-RPE1 H2B-GFP (kindly provided by Prof. David Pellman, USA) cells were maintained in DMEM F12 medium (Sigma, D8437), supplemented with 10% foetal bovine serum (FBS, A5256801, Gibco). Human colorectal carcinoma HCT116 TP53+/+, HCT116 TP53-/-(both kindly provided by Prof. Bert Vogelstein, USA), and HCT116 H2B-GFP (kindly provided by Prof. Zuzana Storchova, Germany) were cultured in McCoy’s 5A medium (Gibco, 36600-021), supplemented with 10% FBS. Human embryonic kidney HEK293T cells (kindly provided by Prof. Masashi Narita, UK) were maintained in DMEM (Gibco, 41966-029) and supplemented with 10% FBS. All cell lines were verified by short tandem repeat (STR) profiling and were regularly tested for mycoplasma contamination. All the cell lines used in this work were cultured in a sterile, humidified incubator at 37 °C with 5% CO2.

### Single-molecule RNA FISH

Cells were grown on 12mm coverslips (ECN 631-1577, VWR, 150 μm thickness) in 12-or 6-well plates, washed briefly with 1x nuclease-free PBS, and fixed with methanol/acetic acid (75%/25%) at room temperature (RT) for 10 minutes (min). Following fixation, the cells were washed with PBS and permeabilised in 70% ethanol for at least 1 hour (h) at 4° C. For knockdown experiments, cells were transfected with siRNAs or LNA gapmers on the day after plating and were fixed 48 hours after transfection. Post-fixation, coverslips were washed with Stellaris® RNA FISH wash buffer A (SMF-WA1-60, LGC Biosearch Technologies), supplemented with 10% deionized formamide (AM9342, Ambion) for 5 min at RT. *C1QTNF1-AS1* FISH exonic probes (labelled with Quasar® 570 dye, Table XX) were added to Stellaris® RNA FISH hybridisation buffer (SMF-HB1-10, LGC Biosearch Technologies), supplemented with 10% deionised formamide at a final concentration of 125 nM. Hybridisation occurred overnight for up to 16 hours at 37° C in a dark, humid chamber. After hybridisation, coverslips were washed with wash buffer A for 30 minutes at 37° C in the dark. This was followed by a 30-min wash with 4,6-diamidino-2-phenylindole (DAPI) in wash buffer A to counterstain the nuclei (5 ng/ml; D9542 Sigma) at 37° C in the dark. The last wash was done with Stellaris® wash buffer B (SMF-WB1-20, LGC Biosearch Technologies) at RT for 5 min. For signal enhancement, coverslips were equilibrated with Base Glucose Buffer composed of 2x saline-sodium citrate (SSC) buffer (10515203 20x UltraPure™ Invitrogen™, Fisher Scientific), 0.4% glucose solution (∼20% in H2O BioUltra, 49163, Sigma-Aldrich) and 20 mM RNase-free Tris pH 8.0 (1 M, 10259194, Fisher Scientific) in RNase-free H_2_O. Then, coverslips were incubated for 5 min in Base Glucose Buffer supplemented with a 1:100 dilution of glucose oxidase (3.7 mg/ml; G7141, Sigma-Aldrich) and catalase (4 mg/ml; C1345, Sigma-Aldrich). Finally, the coverslips were mounted with ProLong™ Gold antifade (P36934, Invitrogen) on a glass slide and left to cure overnight before image acquisition. Z-stacks with 200 nm z-step capturing the entire cell volume were acquired with a GE wide-field DeltaVision Elite microscope (Cytiva) with an Olympus UPlanSApo 100×/1.40-numerical aperture oil immersion objective lens and a PCO Edge sCMOS camera using appropriate filters. The built-in SoftWoRx Imaging software (Applied Precision) was used to deconvolve the three-dimensional stacks. Images were max projected using FIJI ImageJ. RNA FISH signal was counted manually using the Multi point tool in FIJI.

### RNA isolation, cDNA synthesis and qPCR

RNA (1 µg) was extracted from cells using the RNeasy® mini kit (74106, QIAGEN) and treated with DNase I (79254, QIAGEN) according to the manufacturer’s instructions. cDNA was prepared using the QuantiTect® Reverse Transcription Kit (205313, QIAGEN), including an additional wipeout step to eliminate genomic DNA contamination. qPCR was performed on a QuantStudio™ 7 Flex (ThermoFisher Scientific) using PowerUp™ SYBR™ Green Master Mix (A25742, Applied Biosystems). Thermocycling parameters were set to 50 °C for 2 min, 95 °C for 2 min, followed by 40 cycles of 95 °C for 1 sec and 60 °C for 30 sec. Two reference genes (*GAPDH* and *RPS18*) were used to normalise expression levels using the 2−ΔΔCT method. *C1QTNF1/CTRP1* primers for total *C1QTNF1/CTRP1* expression targeting multiple transcript isoforms (NM_198593, NM_030968, NM_153372, XM_006721667, XM_006721665, XM_006721663, XM_006721666, XM_006721664) were supplied by QIAGEN (GeneGlobe ID: QT00044443, Cat. No. 249900, QuantiTect Primer Assays). The sequences of qPCR primers were designed using the web interface Primer3Plus^69^ and are provided in Supplementary Information.

### siRNA and LNA depletion experiments

Cells were transfected at 60-70% confluency with Lipofectamine RNAiMAX (13778150, Invitrogen) following the manufacturer’s instructions. Small interfering RNAs, siRNAs (a pool of four; Horizon Discovery), and antisense Locked Nucleic Acid (LNA) GapmeRs (QIAGEN) were used at final concentrations of 5 nM and 10 nM, respectively. Lipofectamine RNAiMAX and siRNA or LNA gapmer were diluted separately in Opti-MEM™ medium (Gibco, 31985-047). Cells were transfected 24 hrs after plating and harvested for further experimental procedures 24, 48, or 96 hrs after transfection, depending on experimental requirements. Two negative controls, silencer™ Negative Control No. 1 siRNA (Ambion, 4390843) and nontargeting LNA A (300611-00, QIAGEN), were used. For double knockdown experiments, HCT116 cells were plated and transfected the next day with either negative control siRNA (Ctl, from Ambion) and siRNAs targeting RSRC2, in combination with *C1QTNF1-AS1* LNA 1 or Ctl LNA A. siRNAs were combined at 5 nM each to make up a final concentration of 10 nM. For a combination of siRNA with LNA gapmers, siRNAs were used at 5 nM and LNAs at 10 nM, resulting in a final concentration of 15 nM. Cells were always transfected with an equal final concentration of siRNA and LNA gapmers for double knockdown experiments. The list of siRNAs and LNA gapmers is provided in the Supplementary Information.

### Cas13-mediated depletion of *C1QTNF1-AS1*

Guide RNAs (gRNAs) against *C1QTNF1-AS1* were designed using gRNA design tool (https://cas13design.nygenome.org)^37^. Single-stranded gRNA oligonucleotides (Sigma) were 5’-phosphorylated with T4 Polynucleotide Kinase (M0201S, NEB) at 37 °C for 30 min. Forward and reverse oligonucleotides were mixed in a 1:1 molar ratio and annealed at gradually decreasing temperatures from approximately 95-100°C to RT in a glass beaker with the boiling hot water. 5 µg of pxR003 CasRx gRNA cloning backbone (pUC19; 109053, Addgene) was linearised with 10 Units of BbsI restriction enzyme (R3539, NEB) at 37 °C for 4 hrs. BbsI was then heat-inactivated at 65 °C for 20 min. The linearised plasmid was run on 1% agarose gel, and the corresponding band was cut and extracted using QIAquick® Gel Extraction Kit (28704, QIAGEN). Annealed oligonucleotides were ligated into linearised pxR003 with Quick Ligation Kit (M2200S, NEB) for 20 min at RT. NEB5α (C2987H, NEB) cells were transformed with the ligated pxR003 vector according to the manufacturer’s guidelines for transformation. Four bacterial colonies were chosen per gRNA construct and DNA was purified with QIAprep® Spin Miniprep Kit (27104, QIAGEN). All clones were verified by Sanger sequencing. Finally, the HCT116 cells were plated at 80,000 cells/well in 24-well plates for Cas13-mediated depletion of *C1QTNF1-AS1*. 24 hrs after plating, the cells were transfected with the following combination of vectors: pxR003 gRNA vector (200ng) + pxR001 EF1a-CasRx-2A-EGFP (active CasRx, 200ng) (109049, Addgene) using Mirus TransIT-X2® (MIR6004, Mirus Bio). Cells were collected for RNA extraction 48 hrs after transfection and qPCR was performed to verify the *C1QTNF1-AS1* depletion. The list of *C1QTNF1-AS1* guide and negative control sequences is provided in the Supplementary Information.

### Cell proliferation assays

Cells were seeded at 15,000 cells/well (HCT116) on 24-well plates (Corning). The following day, the cells were transfected with *C1QTNF1-AS1* LNAs, including the negative control LNA A. The plates were placed in an IncuCyte® S3 Live-cell Analysis System (Essen BioScience, Ltd; Sartorius Group) 24 hours post-transfection to monitor cell proliferation in real time. The IncuCyte® S3 System was set up to take 4 phase-contrast images per well (4x objective) every 3 hours for up to 3 or 4 days. For analysis, confluence masking was performed on the IncuCyte® S3 Software to obtain an estimate of cell confluence (%).

### Immunofluorescence

The cells were seeded in 12-or 6-well plates on 12mm coverslips (ECN 631-1577, VWR, 150 μm thickness). On the following day, the cells were transfected with siRNAs, Cas13 or LNAs as per the manufacturer’s specifications. After 48 hrs, the coverslips were washed once with 1xPBS and fixed immediately depending on the antibody; methanol, PTEM-F buffer or paraformaldehyde (PFA). *Methanol fixation*: slides were fixed in 99.9% ice-cold methanol (Acros Organics, 167830025) and incubated at −20 °C for 10 min. *PFA fixation*: slides were fixed in 4 % PFA + PBS (28908, Thermo Scientific) and incubated at RT for 15 min. *PTEM-F fixation*: slides were fixed in PTEM-F buffer (20 mM PIPES, 0.2 % Triton X-100, 10 mM EGTA, 1 mM MgCl_2_, 4 % PFA) at RT for 15 min. Slides fixed in methanol or PFA were permeabilised in PBS/0.5 % Tween20/0.5 % Triton X-100, while the slides fixed in PTEM-F were permeabilised in PBS/0.2 % Triton X-100. All permeabilisation took place at RT for 5 min. Slides were blocked in 5 % BSA/ PBS at 4 °C for 1 h, and the primary antibodies (Supplementary Information) were added overnight at 4 °C. Cells were washed 3x for 10 min with PBS/0.1%Tween-20 and then incubated with secondary antibodies diluted in blocking-buffer for 1 h at RT. After washing again 3x for 10 min with PBS/0.1%Tween-20, the cells were stained with Hoescht (1 µg/ml, 33258, Sigma-Aldrich, diluted in PBS) at RT for 10 min. Coverslips were washed with 1xPBS and water before being mounted in ProLong™ Diamond antifade mountant (P36961, Invitrogen).

### Image processing and quantification

Congression defects, nuclear speckle colocalisation and nuclear intensity analysis were performed on slides imaged with a GE widefield DeltaVision Elite High-Resolution Microscope (Cytiva) using an Olympus UPlanSApo 100×1.40-numerical aperture oil immersion objective lens. Analysis of RSRC2, PCNT, CDK5RAP2 and y-tubulin localisation at the centrosomes was imaged with an Eclipse Ti-E inverted microscope (Nikon) equipped with a CSU-X1 Zyla 4.2 camera (Ti-E, Zyla; Andor), including a Yokogawa Spinning Disk, a precision motorised stage, and Nikon Perfect Focus, all controlled by NIS-Elements Software (Nikon) using a CFI Plan Apochromat Lambda D 100× 1.45-NA oil objective (Nikon). For CREST staining, optical slices were taken at 200 nm intervals and 100x oil immersion objective was used. For other antibodies, optical slices were set to 500 nm thickness, and either 100x or 60x oil objective was used.

#### Congression defects

Cells at the metaphase stage of mitosis were imaged using CREST staining to identify misaligned chromosomes. Metaphases were classified as either normal (aligned to the metaphase plate) or congression defects (unaligned chromosomes towards the spindle poles). Congression defects were presented as a percentage of the total metaphase count.

#### Colocalisation of RSRC2 with SC35, a nuclear speckle marker

RSRC2 and SC35 colocalisation analysis was quantified on images stained with both antibodies using the JACoP BIOP plugin for FIJI ImageJ^70^. Sum-projected images were segmented using thresholding to separate the target areas. The colocalisation was reported as the Manders’ coefficient, by measuring the level of overlap between the thresholded pixels from the RSRC2 and SC35 staining. Costes’ randomisation was performed to ensure colocalisation was genuine where we performed 200 rounds of shuffling for the pixels in the measured area. Only data points which met significance following Costes’ randomisation were included in the final analysis.

#### Nuclear RSRC2 intensity

Fluorescence intensity of RSRC2 in the nucleus was analysed using CellProfiler^71^. Images were z-projected using ‘sum slices’ and CellProfiler was used to identify the nuclear area and measure the fluorescence intensity of RSRC2 signal within the nucleus. The following modules were used: *RescaleIntensity* (for both DAPI and RSRC2 channels); *IdentifyPrimaryObjects* based on the DAPI signal, using minimum cross-entropy thresholding to identify nuclei; *FilterObjects*, using AreaShape perimeter, eccentricity and form factor to remove poorly segmented nuclei; *MeasureObjectIntensity* using the filtered nuclei objects to measure fluorescence intensity (Integrated Density) of the RSRC2 channel.

#### Localisation to centrosomes

RSRC2, centrin (Cen 2/3), PCNT, CDK5RAP2 and y-tubulin localisation to centrosomes was analysed in FIJI ImageJ using sum-projected images. A box was drawn around the centrosomes using centrin as a marker, measuring the raw integrated density of RSRC2/PCNT/CDK5RAP2/ y-tubulin and centrin (Cen 2/3). Background signal was accounted for by taking a further measurement close by the centrosome to subtract from the intensity measurements.

### Time-lapse microscopy imaging

HCT116 GFP-H2B (30,000 cells/well) and hTERT-RPE1 GFP-H2B cells (15,000 cells/well) were seeded in 4-compartment glass-bottom cell culture dishes (CELLview™ cell culture dish,627870) with a glass thickness of 175 µm ± 15 µm. Cells were transfected the following day, and live-cell imaging was performed 24-48 hrs, depending on the experiment, after transfection using a GE widefield DeltaVision Elite High-Resolution Microscope (Cytiva). Microscope settings for imaging hTERT-RPE1 GFP-H2B cells were as follows: 10% laser intensity of the FITC (Fluorescein isothiocyanate) channel with a 0.05-second exposure time, 10 optical sections of 2 µm thickness, 4x gain, and 2×2 binning. HCT116 GFP-H2B cells were imaged using similar settings, apart from the exposure time being changed to 0.1 second. Cells were imaged every 3 minutes at 4 or 5 positions per compartment for 10 to 12 hrs. Cells were maintained in a microscope stage incubator at 37 °C and 5% CO2 humidified atmosphere throughout the experiment. The mitotic duration was analysed by visual inspection of the images, counting the time from nuclear envelope breakdown to the onset of anaphase.

### FACS preparation and analysis

Cultured cells were trypsinised and washed with ice-cold PBS, followed by centrifugation. Cells were fixed in 70% ethanol and incubated overnight at 4°C. Fixed cells were pelleted by centrifugation and washed twice in PBS before being resuspended in PBS. RNase A (GE101-01, Generon) was added to a final concentration of 50 µg/ml and the samples were incubated at 37°C for 30 min. The cell suspension was filtered using the 40µm sterile cell strainers, (FisherBrand, 22363547) and incubated with propidium iodide (20 µg/ml, P4170, Sigma Aldrich) on ice in the dark for 30 min. Cells were analysed using a LSR Fortessa (BD Biosciences) and analysed with the FlowJo (v10.8.1).

### Western blot analysis

HCT116 cells were plated at 300,000 cells/dish on 60 mm² dishes (Corning). On the following day, the cells were transfected with siRNA or LNA gapmers, and after 48 hrs, they were trypsinised, washed twice with ice-cold 1x PBS, and pelleted at 1,000 rpm for 3 minutes at 4°C. The pellets were lysed in NP-40 lysis buffer composed of 50 mM Tris-HCl pH 8 (15568025, Invitrogen), 125 mM NaCl (71386, Sigma-Aldrich), 1 % NP-40 (85124, Thermo Scientific), 2 mM EDTA (15575020, Invitrogen), 1 mM PMSF (93482, Sigma-Aldrich), cOmplete™ Mini EDTA-free protease inhibitor cocktail (11836170001, Roche) and phosphatase inhibitors (2 mM sodium fluoride, 201154, Sigma-Aldrich; and 1 mM sodium orthovanadate, S6508, Sigma-Aldrich). The samples were incubated on ice for 25 min and then centrifuged for 3 min at 12,000 x g at 4 °C. The supernatant was collected, and the protein concentration was estimated using the Bio-Rad Protein Assay (5000006, Bio-Rad). Proteins (20 µg) were denatured in 6x SDS buffer (0.375 M Tris pH 6.8, 12 % SDS, 60 % glycerol, 0.6 M DTT, 0.06 % bromophenol blue) at 95 °C for 5 min. The proteins were then separated using Bolt® 4–12 % Bis-Tris Plus Gel (Thermo Fisher Scientific, NW04120BOX) in MOPS buffer (Thermo Fisher Scientific, B0001-02). Precision Plus Protein Standards (161-0373, Bio-Rad) were used as a protein standard. The proteins were then transferred to a nitrocellulose membrane (1620115, Bio-Rad) using 1x Tris-Glycine transfer buffer with 20 % methanol and blocked with 5 % non-fat milk in TBS-T (50 mM Tris, 150 mM NaCl, 0.1 % Tween-20) for 1 hour at RT. The membranes were incubated with primary antibodies in 5 % milk in TBS-T at 4 °C overnight. The following antibodies used were RSRC2 (NBP1-83787, Novus, dilution 1:1000), DIAPH2 (DP4511, ECM Biosciences, dilution 1:1000), CTRP1/C1TNF1 (ab25973, Abcam, dilution 1:500), PCNT (ab4448, Abcam, dilution 1:1000), CDK5RAP2 (702394, Invitrogen, dilution 1:1000), GAPDH (Cell Signaling, 14C10, 2118, dilution 1:1000), and β-tubulin (T019, Sigma, dilution 1:2000). The membranes were washed 3 times the next day with TBS-T and incubated with horseradish peroxidase secondary antibodies (Agilent Dako, P0447 and P0448, dilution 1:2000). Immunobands were detected with SuperSignal™ West Pico PLUS Chemiluminescent Substrate (34580, Thermo Scientific), and the signal was developed using CL-XPosure films (34089, Thermo Scientific). Quantification of immunoblots normalised against appropriate loading controls was performed using ImageJ. Uncropped scans of the immunoblots are provided in Supplementary Figure 8.

### Lentiviral overexpression of *C1QTNF1-AS1* lncRNA

To generate lentivirus, HEK293T cells were plated and transfected with 15 µg of DNA, composed of 9 µg of the lentiviral vector DNA containing the transgene (e.g. lincXpress), 4 µg of psPAX.2 packaging vector (12260, Addgene), and 2 µg of pMD2.G envelope expressing vector (12259, Addgene) in the final transfection volume of 1.5 ml (including 45 µl of Trans-Lt1 transfection reagent, MIR6004, Mirus) using OptiMEM medium (31985070, Gibco). The transfection reaction was incubated at RT for 30 min before adding it to the cells. Twenty-four hours post-transduction, the medium was refreshed. Viral supernatant was collected 48 and 72 hrs post-transduction, spun down at 1,800 × g for 5 min at 4 °C, filtered through a 45 µm filter, and stored in cryovials at −80°C. For overexpression of *C1QTNF1-AS1* in HCT116 cells, *C1QTNF1-AS1* mature RNA sequence (Labomics, based on Gencode vs30) and negative control vector (scrambled *C1QTNF1-AS1* sequence) were cloned into the pLenti6.3/TO/V5-DEST vector (also known as lincXpress; kindly provided by John Rinn, University of Colorado) using the Gateway cloning strategy. HCT116 cells were seeded at 70,000 cells/well on 12-well plates, and the following day, the viral supernatant and appropriate cell medium without antibiotics were added in the presence of polybrene in a dropwise fashion (5 μg/ml, TR100-3, Sigma-Aldrich). Cells were harvested 48 hrs after infection. In the rescue experiment, HCT116 cells were plated at the same density, and the next day, the cells were transfected with control and RSRC2 siRNAs at a final concentration of 10 nM. The following day, cells were transduced with lentiviral particles containing the lincXpress *C1QTNF1-AS1* construct (full) or the negative control scrambled *C1QTNF1-AS1* (scr). Twenty-four hours post-viral infection, cells were harvested for RNA extraction, live cell imaging, or immunofluorescence. The list of *C1QTNF1-AS1* full and scramble sequences is provided in the Supplementary Information.

### In cell protein-RNA interaction (incPRINT)

The incPRINT experiment was performed as described previously^35^. Briefly, the *C1QTNF1-AS1* mature RNA sequence was cloned from the Labomics vector into the 10xMS2 vector (kindly provided by Dr Alena Shkumatava, University of Edinburgh). First, we linearised the 10xMS2 vector with BstBI restriction endonuclease (R0519S, NEB) at 65 °C for 1 h followed by its dephosphorylation using the Quick CIP (M0525S, NEB) at 37 °C for 10 min. The linearised vector was run on a 1% agarose gel, and the products were extracted with the QIAquick Gel Extraction Kit (28704, QIAGEN) according to the manufacturer’s instructions. Secondly, the *C1QTNF1-AS1* sequence (Labomics) was PCR-amplified using *C1QTNF1-AS1* specific primers designed with NEBuilder online software:

*C1QTNF1-AS1 forward (5’-3’):* aaacttaagcttggtaccttcgaaGAAGGAGGAAAGGAGTGAG
*C1QTNF1-AS1 reverse (5’-3’):* cgcgccatcgataccggtttcgaaTTCCAGATCACATATAGAGAG

PCR amplification was performed with Q5 High-Fidelity DNA Polymerase (M0491S, NEB). Amplified insert and linearised 10xMS2 vector were assembled in a 1:3 molar ratio. *C1QTNF1-AS1* insert was cloned upstream of the MS2 stem-loop sequence of the 10xMS2 vector using a Gibson assembly cloning kit (E5510S, NEB) at 50 °C for 15 minutes. The assembled *C1QTNF1-AS1*-10xMS2 vector was transformed into competent E. coli cells NEB5α, and the DNA was purified using the QIAprep® Spin Miniprep Kit. The confirmed positive *C1QTNF1-AS1* 10xMS2 construct was validated with Sanger sequencing. In the *C1QTNF1-AS1* small scale incPRINT experiment, all FLAG-tagged proteins were tested in four replicates. Interaction intensity values (in relative light units, RLU) between *C1QTNF1-AS1*-MS2 and the tested proteins were plotted as the average luminescence across four luciferase replicates. EGFP (enhanced green fluorescent protein) was used as a negative control, and PABPC3 (a polyadenylated RNA binding protein) was included to control for RNA expression. The Xist(C)-MS2 vector were used alongside *C1QTNF1AS1*-MS2 as a positive control for RNA-protein interactions.

### RNA immunoprecipitation (RIP)-qPCR

HCT116 cells were grown on 15 cm^2^ dishes up to ∼90 % confluency. The cells were trypsinised and pelleted at 800 x g for 4 min at 4 °C. The cell pellets were lysed in ice-cold RIP buffer containing 25 mM Tris/HCL pH 7.5, 5 mM EDTA, 0.5 % NP-40, 150 mM KCL (AM9640G, Invitrogen), 0.5 mM DTT supplemented with 100 U/ml RNaseOUT Recombinant Ribonuclease Inhibitor (10777019, Invitrogen), EDTA-free protease inhibitor cocktail (11836170001, Roche) and 1 mM PMSF (93482, Sigma-Aldrich) for 30 min. 1.5 mg of protein total cell lysate was first incubated with 10 μg antibody for 2 hrs at 4 °C (IgG, 2729, Cell Signalling and RSRC2, NBP1-83787, Novus). Pre-washed Dynabeads™ Protein G beads (10003D, Invitrogen) were then added to the antibody-lysate mix for 2 hrs at 4 °C. After incubation, the beads were washed with RIP buffer 3x 10 min at 4 °C on a rotating wheel. RNA was extracted directly from the beads using TRIzol reagent (15596018, Invitrogen) as per the manufacturer’s instructions. Dry RNA pellets were resuspended in 15 μl RNase-free water and subjected to qPCR using the Power SYBR™ Green RNA-to-CT™ 1-Step kit (4389986, Applied Biosystem) as per manufacturer’s guidelines.

### RNA antisense oligonucleotide pulldown coupled with mass spectrometry (MS)

HCT116 cells were plated in 15 cm^2^ dishes and harvested at 90 % confluency. 100 million HCT116 cells were irradiated with 500 mJ/cm^2^ of UV-C (254 nm) in ice-cold 1x PBS on ice. Cells were scraped in ice-cold PBS, spun at 800 × g for 5 min at 4 °C and stored at −80 °C. Amino-C12-LNA-containing oligonucleotides against *C1QTNF1-AS1* and luciferase (negative control) (QIAGEN, custom design) were coupled to Dynabeads™ MyOne™ Carboxylic Acid (65011, Invitrogen) according to the manufacturer’s instructions using 5 nmol LNA oligo per 100 μl beads in the presence of 1-ethyl-3-(3-dimethylaminopropyl)carbodiimide (EDC) (E1769, Sigma-Aldrich) dissolved in MES buffer pH 4.8 (69892, Sigma-Aldrich). Before hybridisation with cell lysate, the oligo-coated beads were equilibrated in RNA pulldown (RP) buffer containing 50 mM Tris/HCl pH 7.5 (15567-027, Gibco), 5 mM EDTA, 500 mM LiCl (L7026, Sigma-Aldrich), 0.5 % DDM (n-Dodecyl β-D-maltoside, D4641, Sigma-Aldrich), 0.2 % SDS (15553027, Invitrogen), 0.1 % Na-deoxycholate (D6750, Sigma-Aldrich), 4 M Urea (U5378, Sigma-Aldrich), 2.5 M TCEP (Tris(2-carboxyethyl)phosphine hydrochloride solution, 646547, Sigma-Aldrich), and protease inhibitors (11836170001, Roche). Then, the oligonucleotide-bead complexes were blocked with RP buffer supplemented with ssDNA 200 μg/ml (salmon sperm DNA, AM9680, Invitrogen), BSA 1 mg/ml (AM2616, Invitrogen) and yeast RNA 200 μg/ml (AM7118, Invitrogen). Cell pellets were lysed in 5 ml RP buffer with murine RNase inhibitor in a 1:500 dilution (M0314L, NEB). Lysates were sonicated in 15 ml tubes with MSE Soniprep 150 Ultrasonicator (5 cycles of 10 sec, with 30 sec intermittent breaks, at setting 15 % at 4 °C) and centrifuged at 16,000 × g for 5 min at 4 °C. At this point, 1 % input was collected. Supernatants were pre-heated to 65 °C with shaking, after which oligo-coated beads were incubated with the lysates for 4 hours at 65 °C with shaking at 1,200 rpm. The beads were washed four times with RP buffer and then three times with 50 mM TEAB buffer (Triethylamonium bicarbonate, 90114, Thermo Scientific) at RT to remove detergents. Beads were resuspended in a final volume of 1 ml 50 mM TEAB buffer from which 50 −100 μl were collected for RNA enrichment analysis. The rest of the buffer (900-950 μl) was discarded, and the beads were stored at −80 °C. Proteins were then subjected to on-bead digestion where the beads were resuspended in 50 mM ABC buffer (ammonium bicarbonate, A6141, Sigma-Aldrich) with 8 M Urea and reduced by adding DTT (Dithiothreitol, 10197777001, Roche) at a final concentration of 10 mM. After a 30 min incubation at RT with 1,200 rpm shaking, samples were alkylated by adding 55 mM iodoacetamide (A3221, Sigma-Aldrich) for 30 min at RT in the dark. Trypsin digestion was performed overnight at RT using 2 μg trypsin (T4799, Sigma-Aldrich) per sample. The next day, samples were desalted using the Stage Tip procedure and recovered in a buffer containing 0.1 % TFA (Trifluoroacetic acid), 0.5 % Acetic Acid and 2 % Acetonitrile for MS analysis. LC-MS analysis was performed on a Q Exactive-plus Orbitrap mass spectrometer coupled with a nanoflow ultimate 3000 RSL nano HPLC platform (Thermo Scientific). The data were analysed using the MaxQuant (version 1.6.3.3) for all mass spectrometry searches^72^. Raw data files were searched against a FASTA file of the Homo sapiens proteome, extracted from Uniprot (2016). Downstream data analyses including data filtering, Log transformation, data normalization, one-sample, or two-sample t-test analysis, category annotation, and data visualizations by scatter or volcano plots, were performed in Perseus software (version 1.6.2.1). Fold change values of median peptide intensities were calculated using three independent oligonucleotides (oligo 1, 3 and 5), and missing values were imputed using the minimal intensity value detected. Evaluation of the MS data was performed to reveal significantly enriched proteins in *C1QTNF1-AS1* pulldown samples over control luciferase pulldown based on a one-sample t-test with Benjamini-Hochberg FDR calculation. To confirm the *C1QTNF1-AS1* enrichment in the oligonucleotide pulldown, RNA from beads from each sample was eluted in 100 μl elution buffer (0.2 % SDS, 2 mM EDTA) at 95 °C for 5 min at 1,200 rpm. Supernatant was collected and mixed with one volume of proteinase K buffer containing 100 mM NaCl, 10 mM Tris/HCl pH 7 (AM9850G, Invitrogen), 1 mM EDTA and 0.5 % SDS with the addition of 1 mg/ml proteinase K (25530049, Invitrogen). The reaction was incubated at 37 °C for 30 min. RNA was extracted with TRIzol™ Reagent (15596018, Invitrogen) and Direct-zol™ RNA Miniprep kit (R2050, Cambridge Bioscience). RNA concentrations were measured using the NanoDrop 2000c UV/IV Spectrophotometer. 1-step RT-qPCR was performed to quantify the RNA enrichment on the beads using Power SYBR™ Green RNA-to-CT™ 1-Step kit (4389986, Applied Biosystems) according to manufacturer’s guidelines where *GAPDH* was used as a housekeeping control.

### RSRC2 Immunoprecipitation (IP) followed by LC-MS/MS-based protein analysis

HCT116 cells (1.4 million) were plated in 10 cm dishes, and the cells were collected at 90 % confluency. In the case of transfection, the same number of cells was plated, and the following day, the cells were transfected with the corresponding LNA gapmers at 25 nm. Four biological replicates were performed for each IP-MS in cells +/-RNase A/T1 Mix or cells treated with Ctl LNA A and *C1QTNF1-AS1* LNA 1 gapmer. The next day, the media was refreshed, and on day four, the cells were washed with ice-cold 1xPBS and lysed in 1 ml of lysis buffer composed of 50 mM Tris-HCl, pH 7.4, 100 mM NaCl, 1 % NP40 detergent, 0.1 % SDS, and 0.5 % Sodium Deoxycholate, supplemented with SUPERase-IN (Invitrogen, AM2694, 40U), phosphatase (Roche PhosStop, 04906837001), and protease inhibitors (Roche Diagnostic GmbH, 14549800), which were directly added to each 10 cm dish. The lysates were transferred into 1.5 ml Eppendorf tubes, cleared with the QIAGEN Shredder (Qiagen, 79656), and spun down at 12,000 g for 10 min at 4 °C. The lysates were then transferred into new tubes, and the concentration was determined with the BCA Assay (Thermo Scientific, 23227). In the meantime, 60 µl of protein G Dynabeads (Dynal, 100.02) were washed 3 times with lysis buffer before the addition of 5 µg of IgG (2729, Cell Signaling) and RSRC2 (NBP1-83787, Novus). The mixture was incubated at RT for 1 h on a rotating wheel. 2 mg of total cell extracts were incubated with 60 µl of protein G Dynabeads coupled to antibodies in a total volume of 1 ml lysis buffer overnight at 4 °C. For the IP-MS sample treated with RNase A/T1 Mix (EN0551, ThermoFisher), 5U of RNase A/T1 Mix was added overnight at 4 °C. RNA was extracted from total cell extracts, and the quality of the RNA was analysed on the TapeStation (Agilent) to demonstrate the efficiency of RNA degradation in the lysates treated with RNase A/T1 Mix. The next day, the beads were washed three times with 1 ml of ice-cold lysis buffer for 5 min at 4 °C on a rotating wheel. To proceed with the IP-MS, the enriched proteins were resuspended with 60 µl of SDS buffer for each sample (2 % SDS, 100 mM Tris pH 7.5, 100 mM DTT). Samples were heated at 95 °C for 5 min, spun down quickly, and by using the magnetic rack, the eluate was transferred into a new Eppendorf tube. 10 µl was used for the western blot to show the efficiency of the IP. The eluates were then subjected to alkylation, detergent removal, and trypsin digestion using the Filter Aided Sample Preparation protocol, followed by desalting using StageTips. Desalted peptides were subsequently lyophilised by vacuum centrifugation, resuspended in 7 μl of A*buffer (2 % acetonitrile, 0.5 % acetic acid and 0.1 % trifluoroacetic acid in water) and analysed on a Q-Exactive plus Orbitrap mass spectrometer (MS) coupled with a nanoflow ultimate 3000 RSL nano HPLC platform (ThermoFisher). The instrument was operated using the Thermo Xcalibur v.4.5 SP1 software. Briefly, 6 μl of each peptide sample was resolved at 250 nl min^−1^ flow rate on an Easy-Spray 50 cm × 75 μm RSLC C18 column (Thermo Fisher), using a 123 min gradient of 3 % to 35 % of buffer B (0.1 % formic acid in acetonitrile) against buffer A (0.1 % formic acid in water). LC-separated samples were infused into the mass spectrometer by electrospray ionization (1.95 kV, 255 °C). The mass spectrometer was operated in data-dependent positive mode, using a TOP15 method in which one MS scan is followed by 15 MS2 scans. The scans were acquired at a range of 375–1,500 *m*/*z*, with a resolution of 70,000 (MS) and 17,500 (MS/MS). A 30-second dynamic exclusion was applied. MaxQuant (version 1.6.3.3) was used for the MS search and protein quantifications^72^ against a FASTAfile of the Homo sapiens proteome extracted from Uniprot (2016). The search for differential phosphorylated peptides (STY) was performed using Maxquant, with phospho (STY) added as a variable modification. All downstream MS data analysis was performed using Perseus software (version 1.6.2.1).

### Splicing analysis

RNA was extracted from control- and RSRC2-siRNA-treated HCT116 cells using the RNeasy kit (74106, QIAGEN) followed by the DNase I (79254, QIAGEN) treatment. cDNA was prepared from 1 ug of total RNA using the QuantiTect Reverse Transcription Kit (205313, QIAGEN), following the manufacturer’s instructions. To analyse the splicing pattern of the alternative events, primers flanking the variable exon (exon 17 within *CENPE*, exon 19 within *PCNT*, exon 19 within *CDK5RAP2*) were designed. *CENPE* transcript isoform containing the exon 17 was amplified by RT-PCR using primers 5ʹ-TCTCTAAGCAGCTCTTACTTTTGA-3ʹ and 5ʹ-CGTGCTGACTATGATAATCTGGT-3ʹ, *PCNT* transcript isoform containing the exon 19 was amplified using primers 5ʹ-ATTTGGCGTGCTTCTTCCAG-3ʹ and 5ʹ-CTCTGCCTGGATGACGCG-3ʹ and CDK5RAP2 isoform containing exon 19 was amplified using primers 5ʹ-TTTTGCAGCCTCCTTTGGAA-3ʹ and 5ʹ-GGGTTTCCAGATAGACTTGCG-3ʹ. PCR reactions were performed using One Taq Hot Start 2x Master Mix with Standard Buffer guideline (M0484S, NEB) as per manufacturer’s instructions. The PCR conditions were the following: *CDK5RAP2*: 94 °C for 30 s, 94 °C 20 s, 53 °C 30 s, 68 °C 1min (repeat 35×), 68 °C 5 min, 4 °C hold; PCNT: 94 °C for 30 s, 94 °C 20 s, 58 °C 30 s, 68 °C 30 s (repeat 35×), 65 °C 5 min, 4 °C hold; *CENPE*: 94 °C for 30 s, 94 °C 20 s, 58 °C 30 s, 68 °C 30 s (repeat 35×), 68 °C 5 min, 4 °C hold. The RT-PCR products were analysed both by normal agarose gel electrophoresis (not shown) and quantified using the QIAxcel DNA High Resolution Kit (1200) (929002, QIAGEN) with the QIAxcel capillary electrophoresis system 100 (9001941, QIAGEN). An alignment marker was injected and run simultaneously with the samples with fragments of 50 bp and 5 kb (929529, QIAGEN). The QIAxcel Size Marker ranging from 50-800 bp (929561, QIAGEN) was used for size and concentration estimation. Samples were analysed using Method OM500. Percentage splicing inclusion (PSI) for each event was calculated by dividing the concentration of the inclusion isoform’s amplicon by the sum of the concentrations of the inclusion and exclusion products.

### RNA library preparation, sequencing and analysis

RNA-seq libraries were prepared from HCT116 cells using the CORALL™ Total RNA-Seq V2 Library Prep Kit with UDIs set B1 (184.96, Lexogen) with Lexogen RiboCop rRNA Depletion Kit for Human/Mouse/Rat V2 following the workflow for generation of long insert sizes. Four biological replicates of cell populations were generated after depleting *C1QTNF1-AS1* or *RSRC2*, with an equal number of replicates for the respective negative controls. The number of PCR amplification cycles was determined by performing a test qPCR using the Lexogen PCR Add-on Kit V2 (020.96, Lexogen) according to the manufacturer’s instructions. Libraries were quantified with Qubit™ dsDNA HS assay kit (Q3285, Invitrogen) and their quality was assessed with the Tapestation (Agilent). Samples were pooled in equimolar ratios and sequenced using 150 bp paired-end reads on Illumina NovaSeq 6000 S4 instrument (Novogene). Each library was sequenced to a depth of >40 million read pairs. The quality of the raw sequencing reads was assessed with FastQC v0.11.9^73^. Extraction of UMIs was performed using the UMI tools v1.1.4^74^ and quality trimming of fastq files using trimgalore v0.6.5. Paired-end reads were aligned to the hg38 build of the human genome using STAR 2.7.0f^75^ and the number of read pairs was counted for each library with rsem v1.3.1 using gencode.v38. Only genes that achieved at least 10 counts in at least four samples were included for downstream analysis. Approximately 80% of uniquely aligned reads were mapped to the human reference genome. Differential gene expression analyses were performed using R package DESeq2^76^, where GRCh38p13.108 was used for gene annotation. For the analyses of alternatively spliced events rMATS v4.1.2^77^ was used with Gencode GRCh38 GTF for alternate exon locations.

## Data and code availability

RNA-sequencing data generated during this study are available in the GEO database (https://www.ncbi.nlm.nih.gov/geo/) under the accession number GSE284007. The mass spectrometry raw files and their associated MaxQuant output files are available in the PRIDE partner repository (http://www.ebi.ac.uk/pride/archive/) under accession numbers: PXD059546 (*C1QTNF1-AS1* oligonucleotide pulldown), PXD059512 (RSRC2 IP-MS in *C1QTNF1-AS1* depleted cells) and PXD059507 (RSRC2 IP-MS with or without RNase).

## Statistical analysis

Graphs and statistical significance were determined using Prism 10 (GraphPad Software), where results are presented as mean ± SEM unless otherwise stated. Statistical analysis was performed on average values for each experiment using: unpaired *t*-test with Welch’s correction, unpaired two-tailed *t*-test, one sample *t*-test for normalised data, Mann-Whitney test, one-way ANOVA with a Dunnett’s multiple comparison test. Normal distribution of all data was assessed using Shapiro-Wilk normality test in Prism before selecting appropriate statistical analysis. *P* values >0.05 were considered statistically not significant. Details of statistical analysis and the number of replicates can be found in the figure and dataset legends.

## Acknowledgements

We are grateful to all the members of the Stojic lab for their comments and discussion of the manuscript. We would like to thank Susana Godinho, Sarah McClelland, and Prabhakar Rajan for their critical reading of the manuscript. We are thankful to the Cancer Research UK (CRUK) Barts Centre Microscopy Facility for their support with image acquisition and CRUK FACS, Proteomics and Bioinformatics Core Facilities at the Barts Cancer Institute (BCI). We thank Kevin Rouault-Pierre and Celine Philippe (BCI, QMUL) for their help with the QIAxcel protocol, John Rinn (University of Colorado) for the lincXpress vector, Maite Huarte and Marta Montes (CIMA, Spain) for their assistance with the oligonucleotide pulldown protocol, and Royal College of Surgeons of England/Devonshire Royal Arch Masons Pump Priming Grant (PO117626 to Prabhakar Rajan, QMUL, UK) for acquisition of the QIAxcel machine. This work was supported by the Barts Charity (MGU0404), Cancer Research UK Career Establishment Award (RCCFEL\100007), Royal Society Research Grant (RGS\R1\231139) and the Academy of Medical Science Springboard Award (SBF006\1026) to L.S. G.G. was supported by an AIRC Fellowship for Abroad. A.N and A. S were supported by the Fondation pour la Recherche Médicale (EQU202003010550) and LABEX DEEP (ANR-11-LABX-0044_DEEP, ANR-10-IDEX-0001-02_PSL) to AS. F.K.M and M.D. were supported by MRC (MR/W001500/2) and BBSRC (BB/X007820/1) project grants to F.K.M. We acknowledge support from the CRUK City of London Centre of Excellence Award Core Funding to Barts Cancer Institute CTRQQR-2021\100004 for access and support from core facilities.

## Disclosure and competing interest’s statement

The authors declare no competing interests.

## Author contributions

L.S. conceived the study. K.G., A.O.C., P.B., G.G., M.D., S.K., A.N., and L.S. performed the experiments. K.G., A.O.C., E.M., A.T., A.N., and L.S. analysed the data. K.G., A.O.C., and L.S. performed the cell biology and biochemical experiments. G.G. performed RNA-seq library preparation. E.M., and A.T. analysed RNA-seq data. K.G., P.B., and M.D. performed proteomics experiments. A.N. performed incPRINT experiment. S.W. helped with image data analysis. J. W. supervised the RNA-seq data analysis. F.K.M. supervised the proteomics data analysis. A.S. supervised the incPRINT data analysis. A.S., and F.K.M. edited the manuscript. L.S. acquired funding, supervised the work and wrote the paper. All authors have read and approved the final version of the manuscript.

## Notes

### Competing Interest Statement

The authors have declared no competing interest.

