## Supplementary material for "The RNA-binding protein RSRC2 promotes mitotic fidelity by interacting with the lncRNA *C1QTNF1-AS1*": Suppl. Information

**The PDF file includes:**

-Supplementary Figures 1-8.
-Lists of primers, LNA gapmers, sequences of siRNA, *C1QTNF1-AS1* RNA FISH sequences, CRISPR/Cas13 guides, LNA pulldown oligos, antibodies for immunofluorescence and Western blotting, and *C1QTNF1-AS1* (Full) and *C1QTNF1-AS1* (Scr) sequences

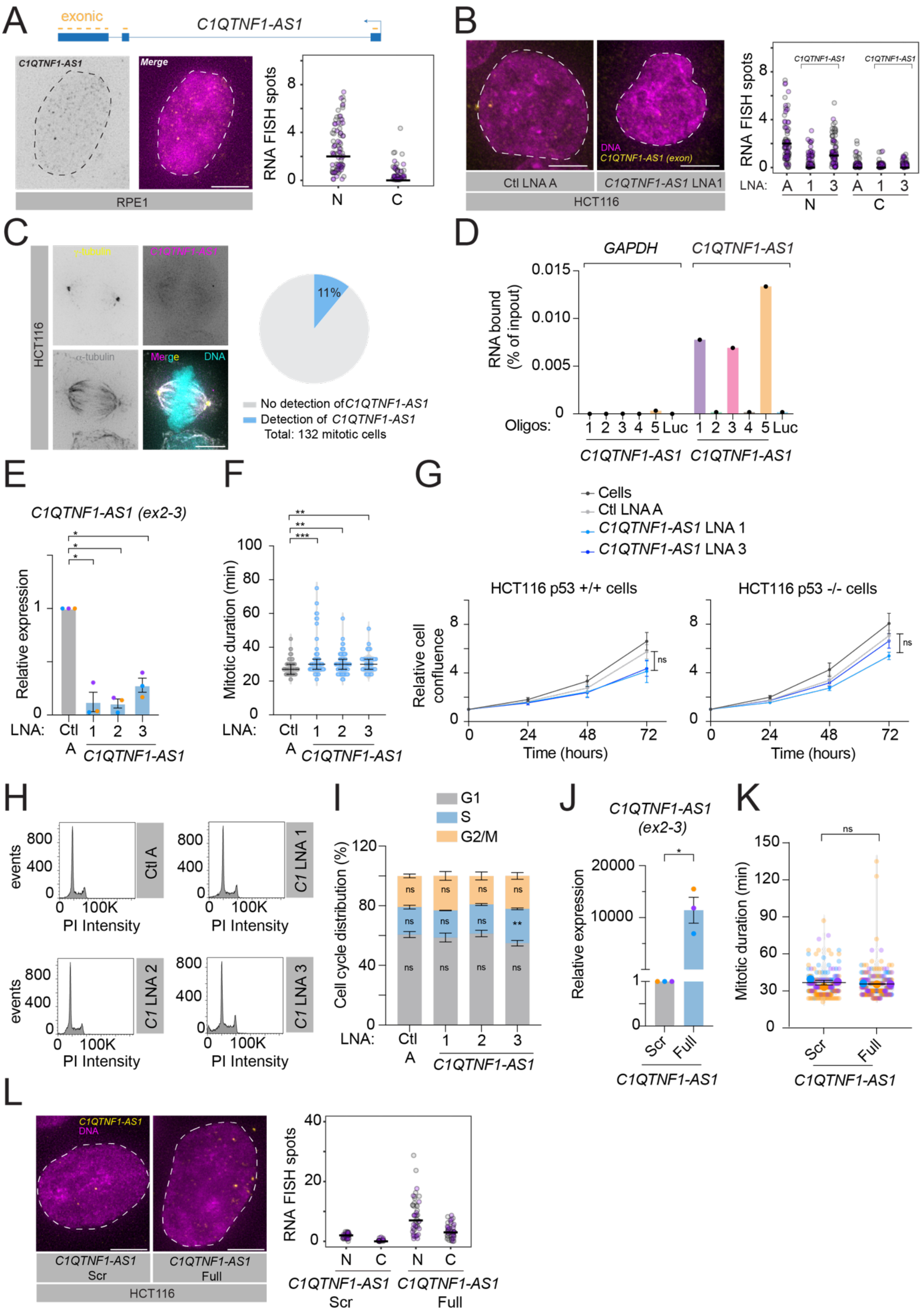


**Supplementary Figure 1. Molecular and cellular characterisation of *C1QTNF1-AS1* in human cells.**

(A) Maximum intensity projections of representative images of *C1QTNF1-AS1* exon smRNA FISH in RPE1 cells showing its nuclear localisation. Nuclei were stained with 4,6-diamidino-2-phenylindole (DAPI, magenta) and outlined with a dashed circle. Right panel: quantification of total transcript in the nucleus (N) and cytoplasm (C), solid line represents the mean. N = 2 ($n_{(RPE1)=}$180).

(B) Maximum intensity projection of representative images of *C1QTNF1-AS1* exon smRNA FISH after LNA-mediated depletion of *C1QTNF1-AS1* in HCT116 cells. Nuclei were stained with DAPI (magenta). Right panel: quantification of total transcript in the nucleus (N) and cytoplasm (C), solid line represents the mean. N = 2 ($n_{(Ctl\ A)}$=112, $n_{(C1QTNF\text{-}AS1\ LNA1)}$=97, $n_{(C1QTNF1\text{-}AS1\ LNA3)}$=97).

(C) Representative image of *C1QTNF1-AS1* exon smRNA FISH (magenta) in HCT116 metaphase cell co-stained for centrosomes ($\gamma$-tubulin, yellow), microtubules ($\alpha$-tubulin, grey), and DNA (Hoescht, cyan). Right panel: quantification of total transcript at mitotic spindle ($n_{(HCT116)}$=132).

(D) qPCR from HCT116 cell extracts showing association of *C1QTNF1-AS1* using ASO 1, 3 and 5 versus Luc. *C1QTNF1-AS1* ASO 2 and 4 did not lead to the enrichment of *C1QTNF1-AS1*. Enrichment of *C1QTNF1-AS1* bound to individual ASO-coupled beads is presented as % of input RNA. *GAPDH* was used as a negative control.

(E) Expression levels of *C1QTNF1-AS1* in RPE1 cells following LNA-mediated depletion of *C1QTNF1-AS1*, as measured by qPCR. Primers spanning mature (ex2-3) *C1QTNF1-AS1* were used. Results are presented relative to negative control (Ctl A) LNA. N = 3.

(F) Quantification of mitotic duration from time-lapse imaging microscopy following LNA-mediated depletion of *C1QTNF1-AS1* in RPE1 H2B-GFP cells. Mitotic duration was measured from nuclear envelope breakdown to anaphase onset. N = 10 ($n_{(Ctl\ A)}$=128, $n_{(C1QTNF1\text{-}AS1\ LNA1)}$=64, $n_{(C1QTNF1\text{-}AS1\ LNA\ 2)}$=106, $n_{(C1QTNF1\text{-}AS1\ LNA3)}$=36).

(G) Quantification of relative cell confluence following LNA-mediated depletion of *C1QTNF1-AS1* in HCT116 p53 +/+ and HCT116 p53-/- cells. N=3.

(H) Representative cell cycle profile following LNA-mediated depletion of *C1QTNF1-AS1* in HCT116 cells. *C1 = C1QTNF1-AS1*.

(I) Quantification of relative cell cycle stages following LNA-mediated depletion of *C1QTNF1-AS1* in HCT116 cells. N = 4.

(J) Expression levels of *C1QTNF1-AS1* following overexpression of *C1QTNF1-AS1* (Scr) and *C1QTNF-AS1*( Full) in HCT116 cells as measured by qPCR. Primers spanning mature (ex2-3) *C1QTNF1-AS1* were used. Results are presented relative to negative control (Scr) plasmid. N = 3.

(K) Quantification of mitotic duration from time-lapse imaging microscopy following overexpression of *C1QTNF1-AS1* (Scr) and *C1QTNF-AS1* (Full) in HCT116 cells. N = 3 ($n_{(C1QTNF1\text{-}AS1\ Scr)}$=237, $n_{(C1QTNF1\text{-}AS1\ Full)}$=256).

(L) Maximum intensity projections of representative images of *C1QTNF1-AS1* exon smRNA FISH in HCT116 cells following overexpression of *C1QTNF1-AS1* (Scr) and *C1QTNF-AS1* (Full) in HCT116 cells. Nuclei were stained with DAPI (magenta) and are outlined with a dashed circle. Right panel: quantification of total transcript in the nucleus (N) and cytoplasm (C), solid line represents the mean. N = 2 ($n_{(C1QTNF1\text{-}AS1\ Scr)}$=76, $n_{(C1QTNF1\text{-}AS1\ Full)}$=92).

Error bars in all panels are shown as mean ± S.E.M. Scale bar, 5 µm. N = number of cells analysed. The following statistics were applied: one-way ANOVA with Dunnett's multiple comparison test in (G) and (I); unpaired t-test with Welch's correction in (E) and (J); Mann-Whitney test in (F) and (K).

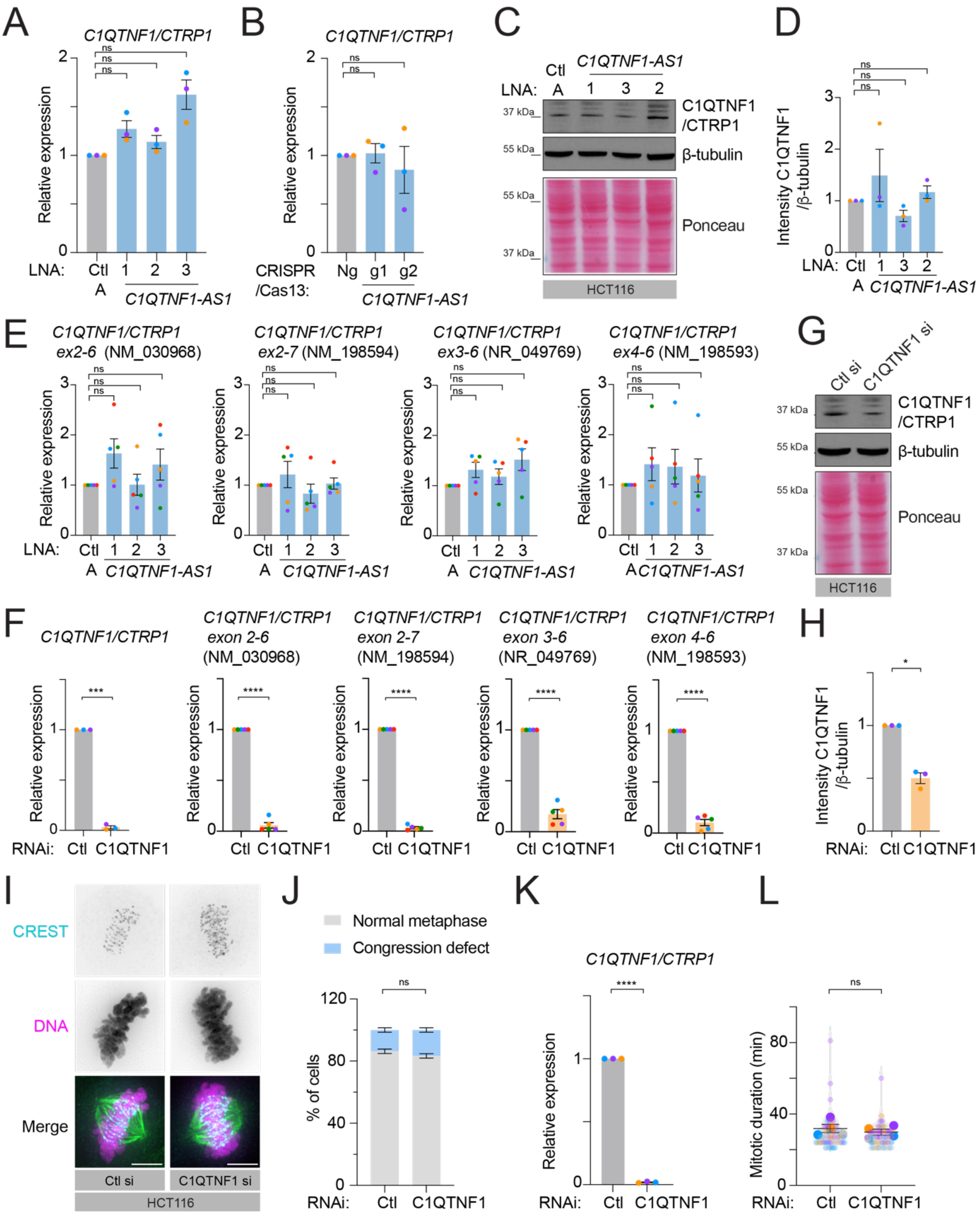


**Supplementary Figure 2. *C1QTNF1-AS1* does not regulate *C1QTNF1/CTRP1* levels and C1QTNF1/CTRP1 is not required for mitosis.**

(A) Expression levels of *C1QTNF1/CTRP1* in HCT116 cells following LNA-mediated depletion of *C1QTNF1-AS1*, as measured by qPCR. Results are presented relative to negative control (Ctl A) LNA. N = 3.

(B) Expression levels of *C1QTNF1/CTRP1* in HCT116 cells following CRISPR/Cas13-mediated depletion of *C1QTNF1-AS1*, as measured by qPCR. Results are presented relative to negative control (Ng) guide. N = 3.

(C) Representative Western blot showing C1QTNF1/CTRP1 protein expression in HCT116 cells following LNA-mediated depletion of *C1QTNF1-AS1*.
(D) Densitometric analysis of C1QTNF1/CTRP1 protein levels from panel (C) relative to control LNA A (Ctl A). N = 3.
(E) Expression levels of *C1QTNF1/CTRP1* isoforms in HCT116 cells following LNA-mediated depletion of *C1QTNF1-AS1*, as measured by qPCR. Primers spanning isoforms NM_030968 (ex2-6), NM_198594 (ex2-7), NM_049769 (ex3-6) and NM_198593 (ex4-6) based on RefSeq annotation were used. Results are presented relative to negative control (Ctl A) LNA. N = 5.
(F) Expression levels of *C1QTNF1/CTRP1* isoforms in HCT116 cells following siRNA-mediated depletion of C1QTNF1/CTRP1, as measured by qPCR. Primers spanning isoforms NM_030968 (ex2-6), NM_198594 (ex2-7), NM_049769 (ex3-6), NM_198593 (ex4-6) and multi-isoform spanning *C1QTNF1/CTRP1* were used. Results are presented relative to negative control (Ctl) siRNA. N = 3 – 5.
(G) Representative Western blot showing C1QTNF1/CTRP1 protein expression in HCT116 cells following siRNA-mediated depletion of C1QTNF1/CTRP1.
(H) Densitometric analysis of C1QTNF1/CTRP1 protein levels from panel (G) relative to control siRNA (Ctl). N = 3.
(I) Representative IF images of metaphase cells stained for kinetochores (CREST, cyan), microtubules ($\alpha$-tubulin, green) and DNA (Hoescht, magenta) after siRNA-mediated depletion of C1QTNF1/CTRP1 in HCT116 cells.
(J) Characterisation of mitotic phenotypes in HCT116 cells after siRNA-mediated depletion of C1QTNF1/CTRP1. The percentage of mitotic cells with normal metaphase plate and congression defect using the antibodies indicated in panel (I). N = 3 ($n_{(Ctl\ si)}$=190, $n_{(C1QTNF1\ si)}$=157).
(K) Expression levels of *C1QTNF1/CTRP1* in RPE1 cells following siRNA-mediated depletion of C1QTNF1/CTRP1, as measured by qPCR. Results are presented relative to negative control (Ctl) siRNA. N = 3.
(L) Quantification of mitotic duration from time-lapse imaging microscopy following siRNA-mediated depletion of C1QTNF1/CTRP1 in RPE1 H2B-GFP cells. Mitotic duration was measured from nuclear envelope breakdown to anaphase onset. N = 4 ($n_{(Ctl\ si)}$=60, $n_{(C1QTNF1\ si)}$=61).

Error bars in all panels are shown as mean ± S.E.M. Scale bar, 5 µm. N = number of cells analysed. An unpaired t-test with Welch's correction was used in panels (A), (B), (D), (E), (F), (H), (J), (K). Mann-Whitney test was used in panel L. β-tubulin and Ponceau staining were used as loading controls for panels (C) and (G).

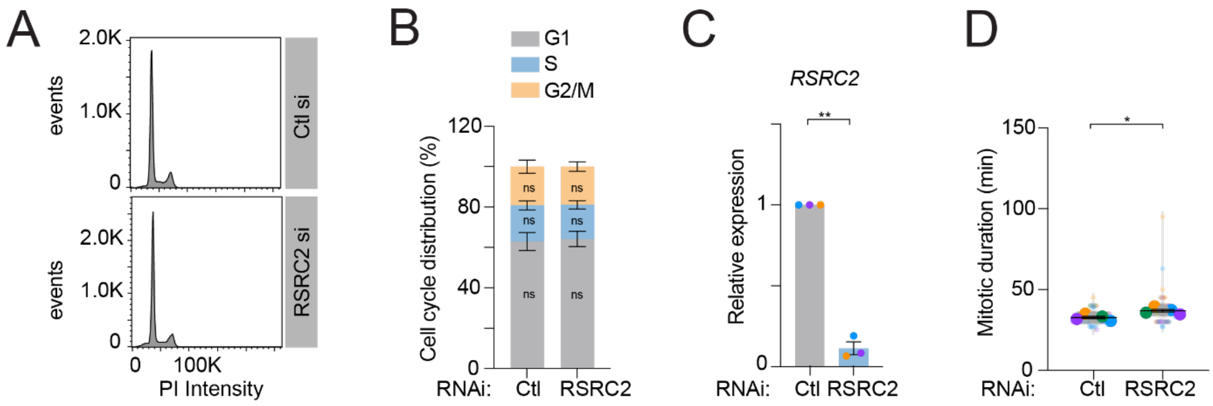


**Supplementary Figure 3. Characterisation of cell cycle progression and mitotic duration in RSRC2-depleted cells.**

(A) Representative cell cycle profile following siRNA-mediated depletion of RSRC2 in HCT116 cells.
(B) Quantification of relative cell cycle stages following siRNA-mediated depletion of RSRC2 in HCT116 cells. N = 4.
(C) Expression levels of *RSRC2* in RPE1 cells following siRNA-mediated depletion of RSRC2, as determined by qPCR. Results are presented relative to negative control siRNA (Ctl). N = 3.
(D) Quantification of mitotic duration from time-lapse imaging microscopy following siRNA-mediated depletion of RSRC2 in RPE1 H2B-GFP cells. N = 5 ($n_{(\text{Ctl si})}$=76; $n_{(\text{RSRC2 si})}$=75).

Error bars in all panels are shown as mean ± S.E.M. The following statistics were applied: unpaired t-test with Welch's correction in panels (B) and (C). Mann-Whitney test was used in panel (D).

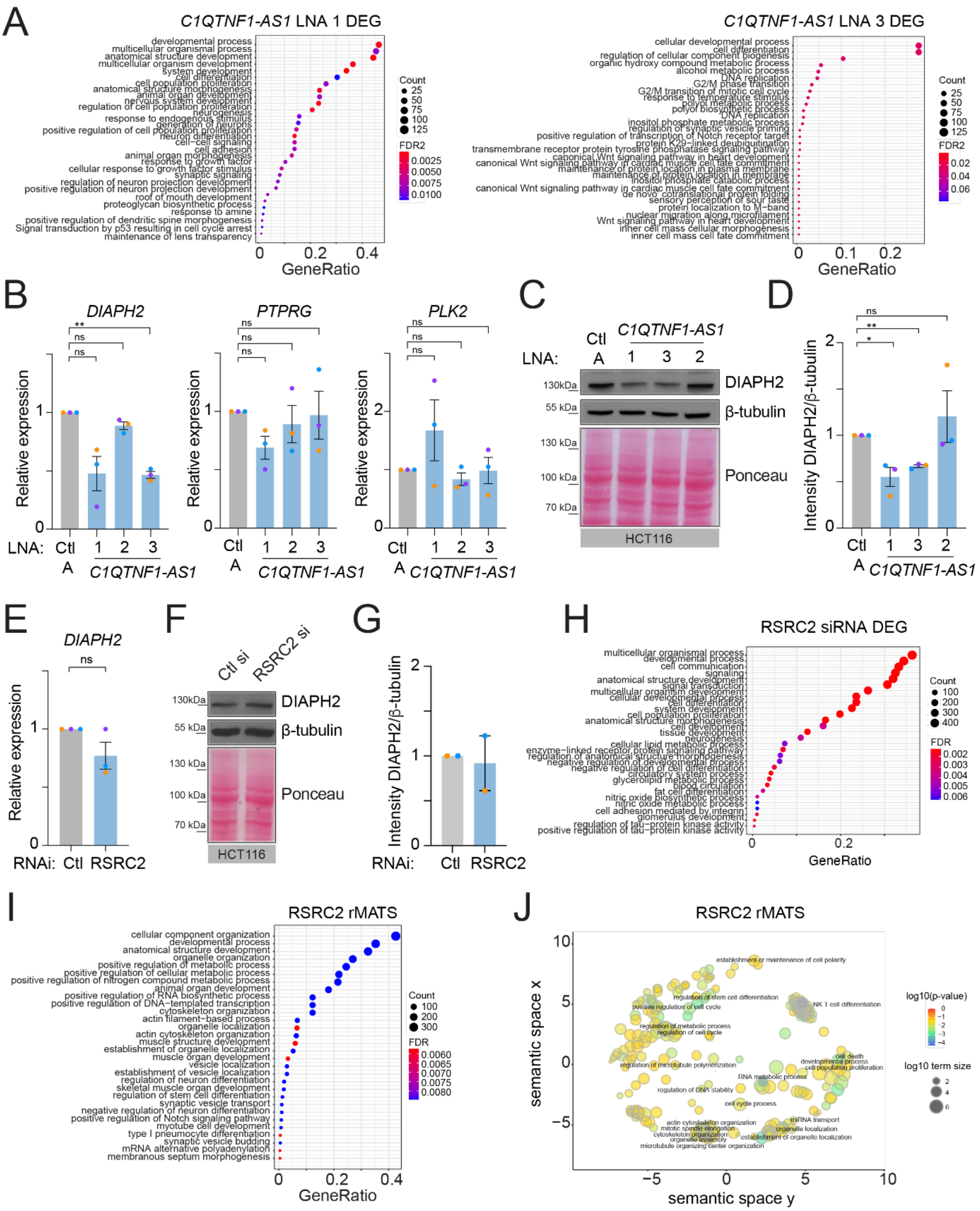


**Supplementary Figure 4. Pathway analysis and validation of DEGs identified in *C1QTNF1-AS1*- and RSRC2-depleted cells by RNA-seq.**

(A) Gene Ontology (GO) analysis of significant differentially expressed genes (DEGs) identified by RNA-seq following LNA-mediated depletion of *C1QTNF1-AS1* in HCT116 cells (as identified in Figure 4A). Top 30 most enriched terms are shown.

(B) Expression levels of *DIAPH2*, *PTPRG* and *PLK2* following LNA-mediated depletion of *C1QTNF1-AS1* in HCT116 cells, measured by qPCR. Results are presented relative to negative control LNA (Ctl A). N = 3.

(C) Representative Western blot showing DIAPH2 protein expression in HCT116 cells following LNA-mediated depletion of *C1QTNF1-AS1*.
(D) Densitometric analysis of DIAPH2 levels from panel (C) relative to control LNA A (Ctl A). N = 3.
(E) Expression levels of *DIAPH2* following siRNA-mediated depletion of RSRC2 in HCT116 cells, measured by qPCR. Results are presented relative to control siRNA (Ctl). N = 3.
(F) Representative Western blot showing DIAPH2 protein expression in HCT116 cells following siRNA-mediated depletion of RSRC2.
(G) Densitometric analysis of DIAPH2 levels from panel (F) relative to control siRNA (Ctl). N = 2.
(H) GO analysis of significant differentially expressed genes identified by RNA-seq following siRNA-mediated depletion of RSRC2 in HCT116 cells (as identified in Figure 4D). Top 30 most enriched terms are shown.
(I) GO analysis of significant differentially spliced genes identified by RNA-seq following siRNA-mediated depletion of RSRC2 in HCT116 cells (as identified in Figure 4H). Top 30 most enriched terms are shown.
(J) GO analysis of significant differentially spliced genes identified by RNA-seq following siRNA-mediated depletion of RSRC2 in HCT116 cells (as identified in Figure 4H). All significant terms are represented, following redundant term reduction performed in Revigo.

Error bars in all panels are shown as mean ± S.E.M. β-tubulin and Ponceau staining were used as loading controls for panels (C) and (F). An unpaired t-test with Welch's correction was used to analyse panels (B), (D) and (E).

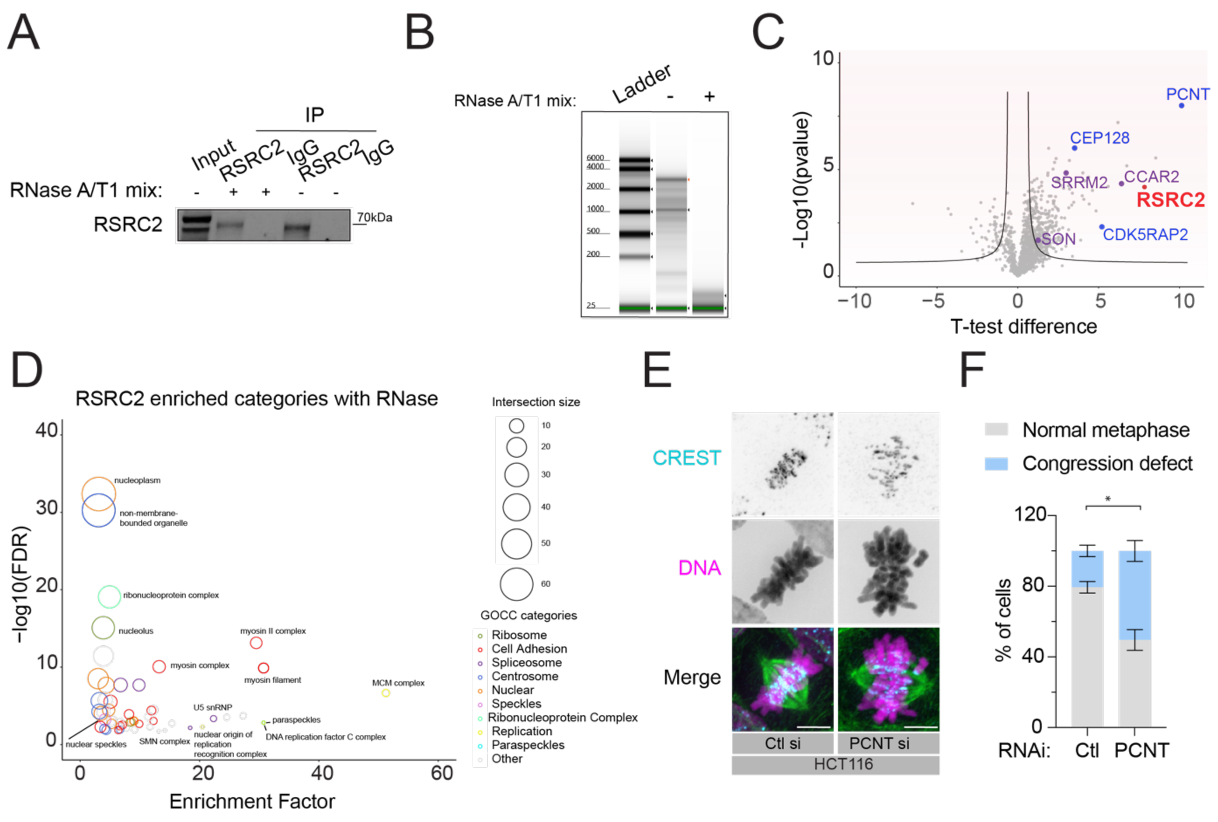


**Supplementary Figure 5. Identification and characterisation of the RNA-dependent RSRC2 interactome.**

(A) Western blot analysis showing the immunoprecipitation (IP) of RSRC2 in HCT116 cells. Pulldown was performed with and without RNaseA/T1 treatment. Rabbit IgG was used as a negative control.
(B) Tapestation analysis showing RNA quality following the IP of RSRC2 from HCT116 cells, with and without RNaseA/T1 treatment.
(C) Volcano plot of PPI for RSRC2 comparing enriched proteins of the RSRC2 IP versus IgG IP in HCT116 cells treated with RNaseA/T1. Curved lines mark the significance boundary (FDR = 1 %). Each dot represents a protein. Selected splicing and centrosome proteins are indicated as some of the top RSRC2 interactors. N = 4. Statistically significant differences were detected using a two-tailed, two-sample t-test with permutation-based FDR.
(D) Fisher's exact test analysis of known protein categories that are over-represented among the RSRC2-interacting proteins shown in panel (C) (Benjamini-Hochberg FDR <0.05). Each circle represents an enriched category from the Gene Ontology Cellular Compartments (GOCC) database, with the circle size representing the number of shared proteins.
(E) Representative IF images of metaphase cells stained for kinetochores (CREST, cyan), microtubules (α-tubulin, green) and DNA (Hoescht, magenta) after siRNA-mediated depletion of PCNT in HCT116 cells. Scale bar, 5 µm.
(F) Characterisation of mitotic phenotypes in HCT116 cells after siRNA-mediated depletion of PCNT. The percentage of mitotic cells with normal metaphase plate and congression defect using the antibodies indicated in panel (E). N = 3 ($n_{(Ctl\ si)}$=337, $n_{(PCNT\ si)}$=355). Error bars are shown as mean ± S.E.M. Unpaired t-test with Welch's correction. N = number of cells analysed.

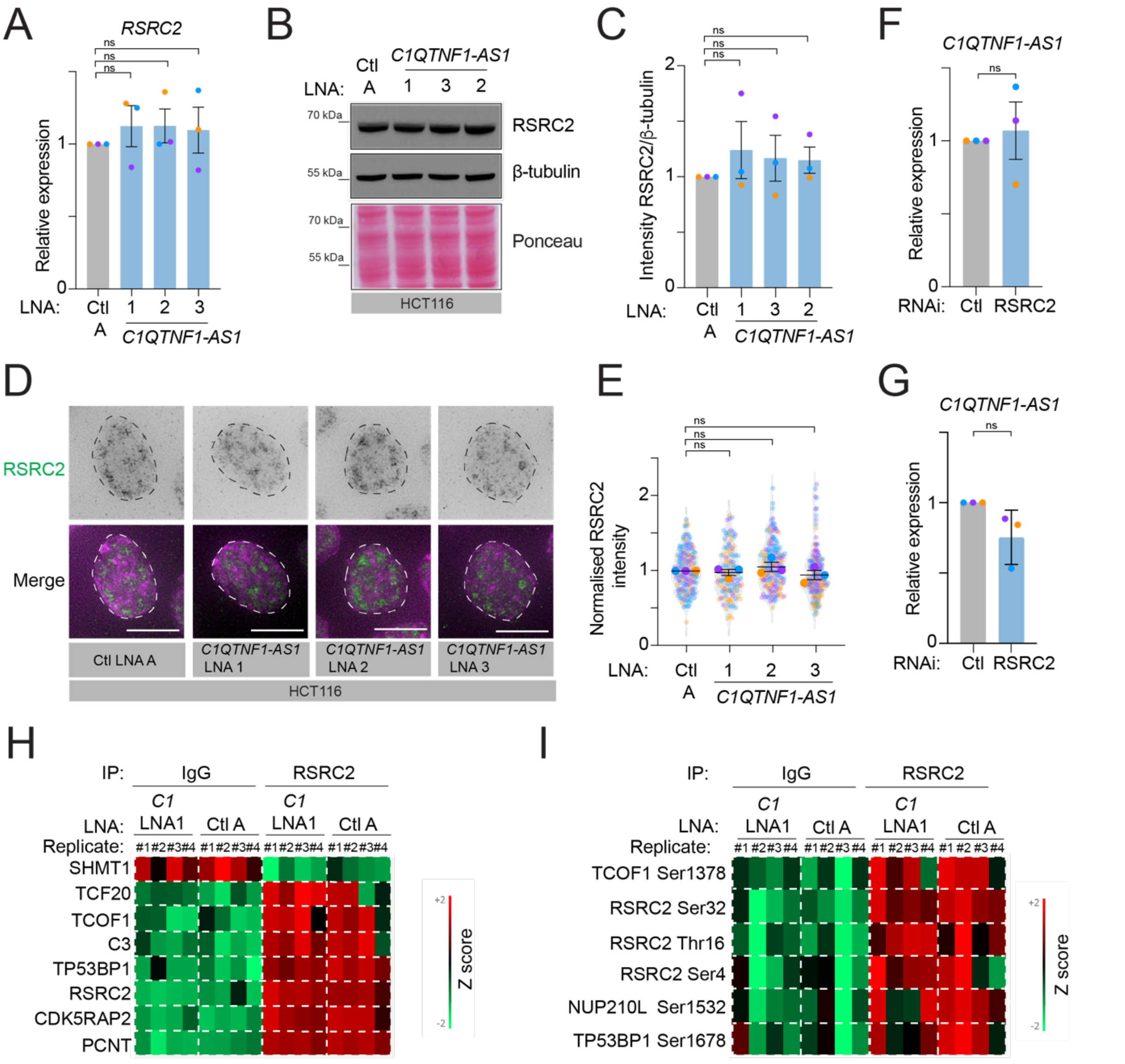


**Supplementary Figure 6. *C1QTNF1-AS1* loss does not affect RSRC2 levels, nuclear localisation, interactome or its phosphorylation status.**

(A) Expression levels of *RSRC2* in HCT116 cells following LNA-mediated depletion of *C1QTNF1-AS1*, measured by qPCR. Results are presented relative to negative control (Ctl A) LNA. N = 3.

(B) Representative Western blot showing RSRC2 total protein levels in HCT116 cells following LNA-mediated depletion of *C1QTNF1-AS1*. β-tubulin and Ponceau staining were used as loading controls.

(C) Densitometric analysis of RSRC2 levels from panel (B) relative to control LNA A (Ctl A). N = 3.

(D) Representative IF images of interphase HCT116 cells stained with RSRC2 (green) and DNA (Hoescht, magenta) after LNA-mediated depletion of *C1QTNF1-AS1*.

(E) Quantification of RSRC2 nuclear intensity in HCT116 cells after LNA-mediated depletion of *C1QTNF1-AS1*. Results are presented relative to control LNA A (Ctl A) treated cells. N = 3 ($n_{(\text{Ctl A})}$=112, $n_{(\textit{C1QTNF1-AS1}\ \text{LNA1})}$=102, $n_{(\textit{C1QTNF1-AS1}\ \text{LNA 2})}$=119, $n_{(\textit{C1QTNF1-AS1}\ \text{LNA3})}$=106).

(F, G) Expression levels of *C1QTNF1-AS1* in HCT116 (F) and RPE1 (G) cells following siRNA-mediated depletion of RSRC2, measured by qPCR. Results are presented relative to negative control siRNA (Ctl). N = 3.

(H) Heatmap showing the top identified RSRC2 interactors in the RSRC2 and IgG IP in HCT116 cells following LNA-mediated depletion of *C1QTNF1-AS1*.

(I) Heatmap showing the RSRC2 phosphorylation status in RSRC2 and IgG IP of HCT116 cells following LNA-mediated depletion of *C1QTNF1-AS1*.

Error bars in all panels are shown as mean ± S.E.M. Scale bar, 5 µm. N = number of cells analysed. An unpaired t-test with Welch's correction was used in panels (A), (C), (F) and (G). One sample t-test (comparing to a hypothetical mean of 1) was used in panel (E). For panels (H) and (I) Z-score of Log2 of LFQ intensity changes for all identified proteins across the different conditions were plotted as a heat map (red-increase; green-decrease). Statistically significant differences were detected using one-way ANOVA, with Benjamini-Hochberg p-value correction for multiple comparison testing, with an FDR cut-off of 0.05 and an S0 of 0.1 (N = 4).

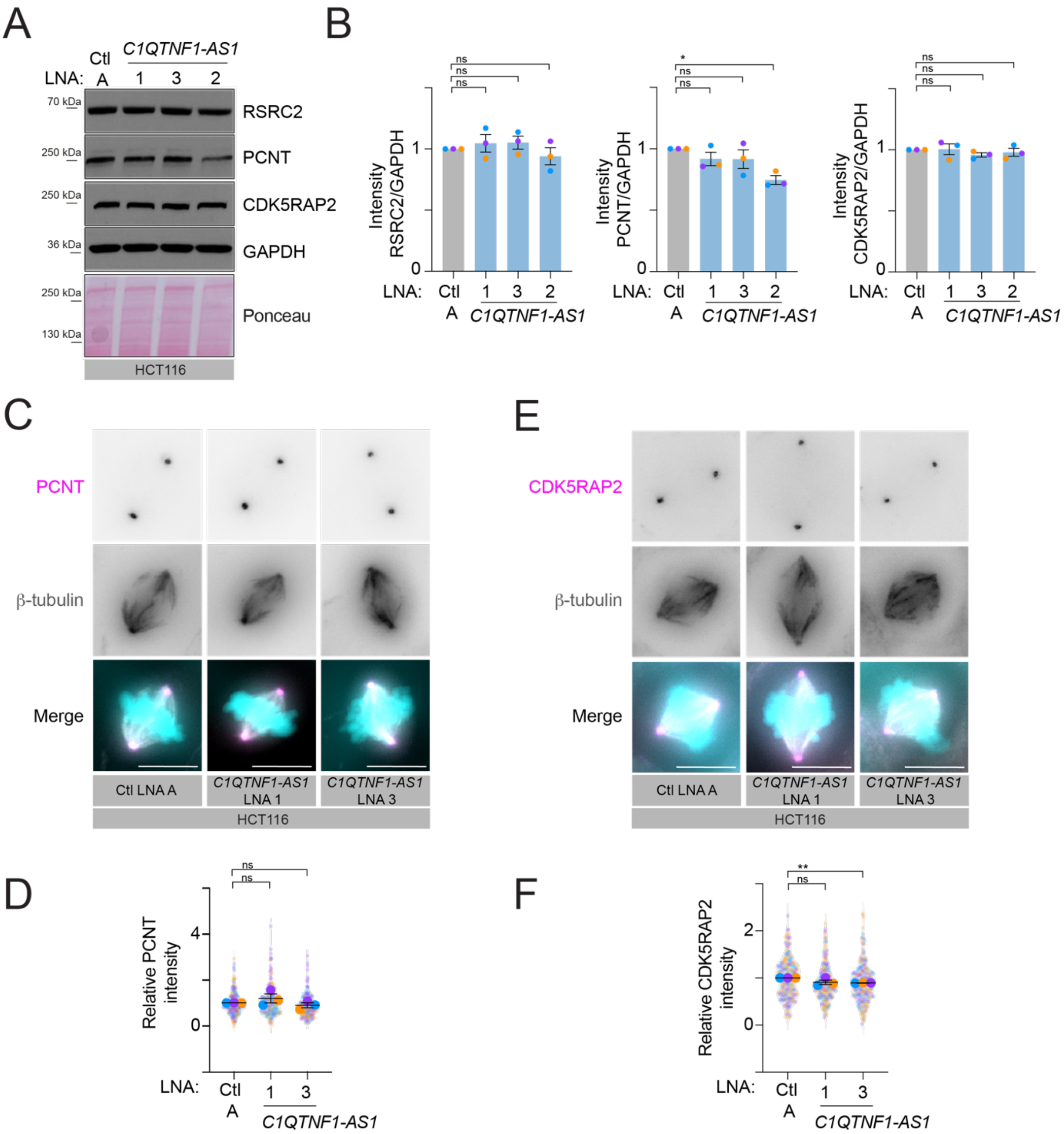


**Supplementary Figure 7. Loss of *C1QTNF1-AS1* does not regulate PCNT and CDK5RAP2 protein levels or their centrosomal localisation.**

(A) Representative Western blot of HCT116 cells probed with RSRC2, PCNT, CDK5RAP2 and GAPDH antibodies following *C1QTNF1-AS1* knockdown. GAPDH and Ponceau staining were used as loading controls.

(B) Densitometric analysis of RSRC2, PCNT and CDK5RAP2 protein levels from panel (A) relative to control LNA A (Ctl A). N = 3.

(C) Representative images of mitotic HCT116 cells stained for PCM (PCNT, magenta), microtubules (β-tubulin, grey) and DNA (Hoescht, cyan) after LNA-mediated depletion of *C1QTNF1-AS1*.

(D) Quantification of PCNT signal intensity around the centrosome of mitotic HCT116 cells for the maximum intensity projections shown in (C). Results are presented relative to control LNA A (Ctl A). N = 3 ($n_{(\text{Ctl A})}$=197, $n_{(\textit{C1QTNF1-AS1}\text{ LNA1})}$=190, $n_{(\textit{C1QTNF1-AS1}\text{ LNA3})}$=190).

(E) Representative images of mitotic HCT116 cells stained for PCM (CDK5RAP2, magenta), microtubules (β-tubulin, grey) and DNA (Hoescht, cyan) after LNA-mediated depletion of *C1QTNF1-AS1*.

(F) Quantification of CDK5RAP2 signal intensity around the centrosome of mitotic HCT116 cells for the maximum intensity projections shown in (E). Results are presented relative to control LNA A (Ctl A). N = 3 ($n_{(Ctl\ A)}$=218, $n_{(C1QTNF1\text{-}AS1\ LNA1)}$=200, $n_{(C1QTNF1\text{-}AS1\ LNA3)}$=216).

Error bars in all panels are shown as mean ± S.E.M. Scale bar, 5 µm. N = number of cells analysed. An unpaired t-test with Welch's correction was used in panel (B). One sample t-test (comparing to a hypothetical mean of 1) was used in panels (D) and (F).

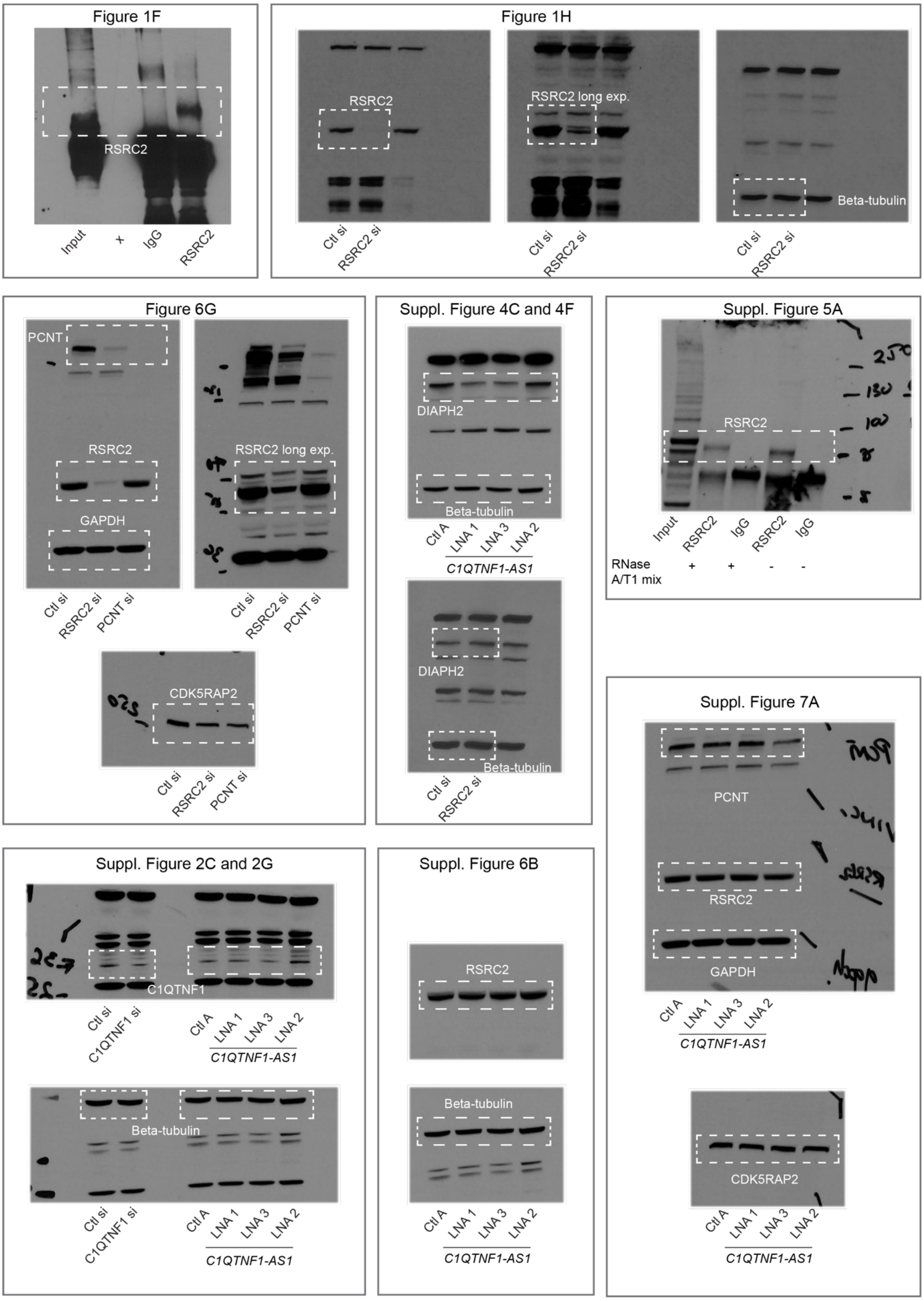


**Supplementary Figure 8. Uncropped pictures of Western blots.**

Regions in white boxes are shown in the main and supplementary figures.

**List of expression primers used for qPCR**

| Target gene | Forward primer (5′ - 3′) | Reverse primer (5′ - 3′) |
|---|---|---|
| *GAPDH* | caacagcctcaagatcatcag | atggactgtggtcatgagtc |
| *RPS18* | atccctgaaaagttccagca | ccctcttggtgaggtcaatg |
| *C1QTNF1-AS1* intron 1.1 | tttggcaagcaagggatgtc | aacactctgcctcttgttgc |
| *C1QTNF1-AS1* exon 2-3 | ggacaggtggagatcaggac | tctccttgtcttctttgcatcc |
| *RSRC2* | ggacaaatcccaatctgctg | gggtttgtgatcttgccatt |
| *PTPRG* | cttgaccttcgtgtgcctcatc | gaaggcacttcacggaaacgtc |
| *PLK2* | caacaatggtgctcacatgagcc | ggagcatctgttgctgggaaaac |
| *DIAPH2* | aaacctgaagtgtccatgaagag | tgcttctgcgttcttttgaact |
| *C1QTNF1/CTRP1 exon 2-6* | ttccttcaccgagtctgtgc | cagcaagagtccctgtccac |
| *C1QTNF1/CTRP1 exon 2-7* | tctgtttccttcaccgagtctg | cctgactgggcctgtatttttc |
| *C1QTNF1/CTRP1 exon 3-6* | tcatcatgatgtcagccacagg | ccagaggcaaaggcaaggag |
| *C1QTNF1/CTRP1 exon 4-6* | cagggcctgggcaaaaatttac | cagcaagagtccctgtccac |
| *C1QTNF1* | QIAGEN GeneGlobe ID: QT00044443, Cat. No. 249900, QuantiTect Primer Assays, Qiagen | |

**List of LNA gapmer sequences**

| Target gene | Sequence 5′ - 3′ | Product number |
|---|---|---|
| Negative control LNA A, QIAGEN | AACACGTCTATACGC | 300611-00 |
| *C1QTNF1-AS1,* LNA gapmer 1 (exon 3), QIAGEN | TACGGCAATGCTTCAG | 300603-00 |
| | | Design ID:591242-1 |
| *C1QTNF1-AS1,* LNA gapmer 2 (intron 1), QIAGEN | GAAGTGAAGTTGATGA | 339511 |
| | | LG00782377 |
| *C1QTNF1-AS1,* LNA gapmer 3 (intron 1), QIAGEN | GACGGACACCAGGAAG | 339511 |
| | | LG00782376 |

**List of siRNA sequences**

| List of siRNA sequences | Sequences (antisense) | Catalogue number |
|---|---|---|
| Negative control siRNA No.1 (Thermo Fisher Scientific, Silence select, Ambion) | / | 4390843 |
| siGENOME Human C1QTNF1 SMART pool, Horizon Discovery | GCACAGCAACCACUACUAC | M-014686-00-0005 |
| | GAACCUCUACGACCACUUC | |
| | GAUCAACAUCACUAUCUUG | |
| | GAGCUGGACACCUACAUCA | |
| siGENOME Human RSRC2 SMART pool, Horizon Discovery | GGAAAUUGAUGGGUAUUAA | M-016134-01-0005 |
| | AGGAAGAAGUAUUUCGAAA | |
| | CCAUUAAACUUGACAGGAC | |
| | AAUUACAAGAACAGCGAGA | |
| siGENOME Human PCNT SMART pool, Horizon Discovery | GCAAAGAACUGUCUGCAAA | M-012172-01-0005 |
| | GAUGAAAGGUGACUUAGAA | |
| | GUAAAGAGAUCACCGAGAA | |
| | GGACGAAGCUUGCUCACUU | |

**List of *C1QTNF1-AS1* smRNA FISH probes (Stellaris)**

| Probe | Sequence 5′ - 3′ |
|---|---|
| C1QTNF-AS1_E_Q570_1 | tgcagcctctagttctct |
| C1QTNF-AS1_E_Q570_2 | cgcagtgagatgctccaa |
| C1QTNF-AS1_E_Q570_3 | caggagcaggacatgctc |
| C1QTNF-AS1_E_Q570_4 | tctatgctggggtccttg |
| C1QTNF-AS1_E_Q570_5 | tctcatgagccagggatc |
| C1QTNF-AS1_E_Q570_6 | caggctctcttggggcag |
| C1QTNF-AS1_E_Q570_7 | gctcaacaatgtgtttcc |
| C1QTNF-AS1_E_Q570_8 | gaaacagcgacgtcagct |
| C1QTNF-AS1_E_Q570_9 | cccactttagagagcagt |
| C1QTNF-AS1_E_Q570_10 | cagtgggacgtcaccatc |
| C1QTNF-AS1_E_Q570_11 | catcttagcagcttctcc |
| C1QTNF-AS1_E_Q570_12 | tctgccaagtttccactt |
| C1QTNF-AS1_E_Q570_13 | caggagccacgagcatgg |
| C1QTNF-AS1_E_Q570_14 | ccaagctgctcagaacgt |
| C1QTNF-AS1_E_Q570_15 | agtcctgatctccacctg |
| C1QTNF-AS1_E_Q570_16 | ggtgtgtccttgcagctg |
| C1QTNF-AS1_E_Q570_17 | ctctttagtttctgtggc |
| C1QTNF-AS1_E_Q570_18 | cctcagagacttcaccat |
| C1QTNF-AS1_E_Q570_19 | tctccttgtcttctttgc |
| C1QTNF-AS1_E_Q570_20 | tcaggctttcattggatt |
| C1QTNF-AS1_E_Q570_21 | tcactttccttaaggtca |

**List of CRISPR/Cas13 guides**

| Target | Sequence |
|---|---|
| Non-targeting guide 1, NT1 | aaacTCACCAGAAGCGTACCATACTC |
| *C1QTNF1-AS1* guide 1 (exon 3) | aaacATTTGCATTCTGTGAGCGAACAC |
| *C1QTNF1-AS1* guide 2 (exon3) | aaacTAGAAGAAAGATTAGGGGGTCAA |

**List of LNA oligonucleotides for RNA pulldown**

| Probe | Sequence |
|---|---|
| *C1QTNF1-AS1* oligo 1, exon 3 | /5AmMC12/ttatactacggcaatgcttca |
| *C1QTNF1-AS1* oligo 2, exon 3 | /5AmMC12/tgcattctgtgagcgaacaca |
| *C1QTNF1-AS1* oligo 3, exon 3 | /5AmMC12/agacttcaccatagactcctt |
| *C1QTNF1-AS1* oligo 4, exon 1 | /5AmMC12/agcgacgtcagctcagctcaa |
| *C1QTNF1-AS1* oligo 5, exon 1 | /5AmMC12/aggacatgctcactccttt |
| Luciferase oligo 1 | /5AmMC12/tcactgcatacgacgattct |
| Luciferase oligo 2 | /5AmMC12/attgggagcttttttgcac |
| Luciferase oligo 3 | /5AmMC12/aaccgggaggtagatgagat |
| Luciferase oligo 4 | /5AmMC12/agttcgcctctttgattaac |
| Luciferase oligo 5 | /5AmMC12/gacgtaatccacgatctctt |

**List of antibodies used for immunofluorescence (IF) and Western blotting**

| **Antibody** | **Supplier** | **Dilution used for IF** | **Fixation method for IF** | **Dilution used for Western blot** |
|---|---|---|---|---|
| CREST | 15-234-0001, Antibodies Inc | 1-1000 | PTEM-F | - |
| α-tubulin (DM1A) | T9026, Sigma | 1-1000 | PTEM-F | - |
| α-tubulin (Rat) | MCA78G, AbD Serotec | 1-500 | PTEM-F | - |
| RSRC2 | NBP1-83787, Novus Biologicals | 1-100 | Methanol/PFA (as indicated) | 1-2000 |
| SC-35 | S4045, Sigma-Aldrich | 1-2000 | PFA | - |
| Centrin-2/3 (Cen2/3) | 04-1624, Sigma-Aldrich | 1-1000 | Methanol | - |
| β-tubulin | T0198, Sigma-Aldrich | - | - | 1-4000 |
| β-tubulin, scFv-S11B, conjugated with Human IgG1 | AA344, ABCD Antibodies | 1-800 | Methanol | - |

| | | | | |
|---|---|---|---|---|
| Pericentrin (PCNT) | ab4448, Abcam | 1-1000 | Methanol | 1-2000 |
| CDK5RAP2 | 702394, Invitrogen | 1-250 | Methanol | 1-5000 |
| C1QTNF1/CTRP1 | ab25973, Abcam | - | - | 1-500 |
| DIAPH2/mDia3 | DP4511, ECM Biosciences | - | - | 1-1000 |
| γ-tubulin | T5192, Sigma-Aldrich | 1-1000 | Methanol | - |
| GAPDH | 2118, Cell Signalling Technology | - | - | 1-5000 |
| Alexa Fluor® 647 goat anti-human | A21445, Invitrogen | 1-1000 | - | - |
| Alexa Fluor® 555 donkey anti-mouse | A31570, Invitrogen | 1-1000 | - | - |
| Alexa Fluor® 488 donkey anti-rabbit | A21206, Invitrogen | 1-1000 | - | - |
| Alexa Fluor® 488 donkey anti-mouse | A21202, Invitrogen | 1-1000 | - | - |
| Alexa Fluor® 555 goat anti-rat | A-21434, Invitrogen | 1-1000 | - | - |
| Amersham ECL Mouse IgG | NA931V, GE Healthcare | - | - | 1-2000 |
| Amersham ECL Rabbit IgG | NA934V, GE Healthcare | - | - | 1-2000 |

**List of target sequences for Lincexpress vectors**

| Target | Sequence |
|---|---|
| *C1QTNF1-AS1*<br>Full (based on Gencode 30) | gaaggaggaaaggagtgagcatgtcctgctcctgcatgtccctgcttaagctcagg<br>actggcccttccaggccaaggaccccagcatagaccccaggacagggccccaa<br>ggatccctggctcatgagagcggcttgctgggctgccccaagagagcctgaagg<br>aaacacattgttgagctgagctgacgtcgctgtttcttccagactgctctctaaagtgg<br>gcagggtagcgaccggccggctccgatggtgacgtcccactgccaaggggtggg<br>agtggggagagtctccacagagcttcggagaagctgctaagatggaaaagtgga<br>aacttggcagacagatccagcctccctggccactggcccatgctcgtggctcctgg<br>atggcgctgccacgttctgagcagcttgggacaggtggagatcaggactggcagc<br>tgcaaggacacaccagagccacagaaactaaagagaatttccaaaaggagtct<br>atggtgaagtctctgaggatgcaaagaagacaaggagaatgaaaatccaatgaa<br>agcctgattgtatttgttgaccttaaggaaagtgattttatggtacagcctctctggaag<br>ggagggtgtgttcgctcacagaatgcaaatacccttgacccccctaatctttcttctag<br>gagtttctcctacagataaacttagaagggtgctcaaataagtaagttcaaggatat<br>cctctgaagcattgccgtagtataaaaaaagcacagataccctcaaaggacatca<br>ttaggggcctggtaaaataaattccacacagtggaacaccgtgtagctctttagaga<br>ataaacagctctctatatgtgatctggaa |
| *C1QTNF1-AS1*<br>Scramble (Scr) | gtggcggatgttggcttagaatcaggattgtgcagatattgaaccttgttgcctcccac<br>cctgcacagcttgatgacccctaaaagtagcagagcttcacaagtactattgggca<br>atcgctccggcgatggtgaaaaaacctactcttcaacacggtcgagcttggtctacc<br>ctcttcagcgaacattacaagaaacagaactcataggcatcaaatgtttgaggacc<br>atggagggatatgcctagacgcgcggtagcaggcaagatcagggtttaacatgg<br>cctcttgaaacttagaggtacagccgccgcgcgtccgcggtagtgcactgcggaa<br>accgcataatatcgtgaggtaggtgggcgaagtggtcttggtgcagcttgaccttaa<br>gtgttaggacccatgtagatagcaaaagcagggcggcggcgtcataaagggttc<br>agcgcacccatgcgaggaagttaggagagaaccaaggacaatcgggccatca<br>ggatgacaggcagaaacgaagtacacacagaagcgatccgtttattaacgcaac<br>attggcggctatcaacacattcgtcctattgtgtaggttgcatggtaattctgtcatataa<br>ggcgtgtccccaacacaataaaggttcgccgctaagctcgccaaatccgatcagg<br>caacaaagtcggatctatactagtctaaagaggcggagaggctagagcttcaatc<br>cgccaccgcgaccaattctaagttaggttagtgtcgactgtgaaagcaaagaccg<br>gcctcaacgggtagtgagcggggatggactggagtcctaagcacacgctgttgcg<br>ctagagaatctaggactcggttactcatcc |
